# CREB: Consistent Reference External Batch Harmonization

**DOI:** 10.64898/2026.03.10.710874

**Authors:** Ameya Kharade, Yiyan Pan, Carmen Andreescu, Helmet Karim

**Affiliations:** Department of Psychiatry, University of Pittsburgh, Pittsburgh, PA; Department of Bioengineering, University of Pittsburgh, Pittsburgh, PA; Department of Psychiatry, Department of Bioengineering, University of Pittsburgh, Pittsburgh, PA; Departments of Psychiatry, Department of Bioengineering, University of Pittsburgh, Pittsburgh, PA

**Keywords:** fMRI Harmonization, Machine Learning, Empirical Bayes, ComBat, Resting State, Connectivity

## Abstract

Machine learning models using functional magnetic resonance imaging (fMRI) are becoming increasingly popular - these models often rely on training data from multiple, large, and publicly available datasets. It is often necessary to harmonize these data across sites and sequences, and algorithms like ComBat are frequently applied to correct for these differences. This has been shown to improve model performance and generalizability. However, applying traditional ComBat necessitates harmonizing all data (train, validation, test, and other unseen external test sets) simultaneously, which leads to potential data leakage and limits application to new unseen data. We introduce Consistent Reference External Batch (CREB) harmonization, a novel extension of ComBat that learns the prior distribution of site effects exclusively from a designated training set. This learned prior serves as a consistent, easily deployable reference point that employs the empirical Bayes framework to update the site effect for any new, external unseen data. This approach enables training, validation, and test sets to be harmonized separately, thereby preventing data leakage, ensuring the integrity of downstream analyses, and application to new unseen data. CREB is different from traditional ComBat in which each sites’ prior distribution is estimated at once, but this cannot be applied to unseen data or data from sites not included in the original set of data. We tested CREB with train data from 2846 participants (ages 18-97 years) across 9 different studies and test data from 1113 participants (ages 18-88 years) from 3 studies. We evaluated the performance of harmonization with functional connectivity and gray matter volume. We show that CREB can effectively harmonize the test data to the train data, and have comparable performance to ComBat. CREB is able to conduct this harmonization in a two-step procedure that prevents leakage and is deployable to new unseen data. Finally, we tested whether CREB could similarly preserve biological variance (e.g., whether age associations were preserved after harmonization). We found that CREB, like ComBat could preserve age associations with both functional connectivity and gray matter volume measures. CREB provides an easily deployable, robust harmonization method to standardize data to a common reference distribution, making it uniquely suitable for training generalizable machine learning models.

## 1 Introduction

Neuroimaging research is evolving rapidly with substantial growth in size and scale of available datasets. This expansion is driven by collaborative efforts between institutions and large open-source datasets collected at multiple facilities with varying sequences. While this increase in data significantly boosts the power of analyses, it simultaneously introduces unique challenges associated with differences in sites. These differences are due to effects related to different scanners, field strengths, acquisition protocols, and manufacturers [Jovicich et al., 2006, Takao et al., 2011]. This is a well-known problem that introduces non-biological signal related artifacts to neuroimaging data. Harmonization is necessary for multi-site data to remove the site variability while preserving biological signal. It is often done with tools like ComBat [Fortin et al., 2018]. ComBat works by assuming that the signal is composed of biological features and non-biological site effects that have both additive and multiplicative effects. Methods like NeuroHarmonize [Pomponio et al., 2020] extend on ComBat by allowing for non-linear modeling of biological signals. This unwanted non-biological variability has been shown to be effectively removed using such tools in diffusion weighted imaging [Fortin et al., 2017], structural imaging [Fortin et al., 2018], and functional imaging [Bostami et al., 2022, Yu et al., 2018]. These tools have significantly improved our ability to analyze data with multiple sites and account for differences between these sites, however there are some major unaddressed challenges. Current tools require harmonization to occur with all data available, however in machine learning, we often want to train on data and apply these models to unseen test data. This results in a second issue that if we harmonize train and test data jointly, we can inadvertently introduce data leakage that can inflate performance [Rosenblatt et al., 2024]. Data leakage is when information about the test data is introduced or exposed to the model during training. Harmonizing train and test sets jointly introduces leakage.

Jointly harmonizing train and test data can cause data leakage between train and test sets that may inflate model performance and confidence [Rosenblatt et al., 2024]. When we deploy machine learning models, we often need to make predictions on new unseen data, which means we need to be able to harmonize this data to the training set. This can only be done if the training data is deployed with the model, which is often not possible due to limitations on data sharing and size of the training data. We have developed a novel extension on ComBat for harmonizing data (including functional and structural imaging measures) that can harmonize train and test data separately, which avoids data leakage, and can be deployed easily without having to upload training data. A key innovation is the estimation of a prior distribution on site effects, which is learned exclusively from this designated training set. This learned prior is then leveraged within an empirical Bayes framework to update the posterior for each new site individually. This approach enables training, validation, and test sets to be harmonized completely independently of one another. In addition, this prior distribution can be deployed with a small ‘bundle’ of size around 13MB that includes simple global statistics needed to approximate this distribution. This prevents data leakage, allows harmonization of unseen data, and is easy to deploy alongside a machine learning model.

There are two major challenges that current approaches face, and they stem from the fact that harmonization approaches require all data to be available to conduct the harmonization. Thus, both NeuroComBat and NeuroHarmonize are fit on all data in a single step. The two challenges that arise are: (1) in machine learning, we typically separate the train and test sets to prevent data leakage - conducting harmonization would potentially result in leakage if the train/test sets are harmonized together [Rosenblatt et al., 2024]. One thought is to harmonize the train data first, train the model, and then harmonize train/test together but this may result in some performance differences. This leads us to another challenge: (2) harmonizing unseen data or new data would require deploying the training data. Machine learning models would be required to deploy the whole dataset to harmonize the new unseen data. In addition, that model may need to be retrained due to loss in performance. Thus, we sought to build on these approaches that resolved these problems of non-incremental harmonization, data leakage, and lack of an easily created and distributable reference point. We propose CREB: Consistent Reference External Batch Harmonization - a two-stage approach (1) CREB Learn - that estimates a site prior in the training set as an easy to store ‘bundle’ of statistics (less than 13MB) and (2) CREB Apply - uses this prior distribution to update the posterior for new unseen sites and harmonizes data to the training data without leakage or need to share the training data. In CREB Learn, the training bundle is created from a collection of large and diverse training datasets covering the whole adult life span, and the priors are estimated across all training sites and features. In CREB Apply, for all sites we want to harmonize, we can use the same prior created on training data. By having a single reference point, data from different sites, especially unseen data, can be harmonized relative to the train data. There have been some approaches in recent years to build harmonization methods that address certain key limitations, such as DC-Combat [Bostami et al., 2022], which allows for decentralized harmonization through COINSTAC but still needs complex orchestration, repeated federation to add new sites, and offers no built-in leakage control. More recently, one group developed ComBat-Predict [Xin et al., 2025] that extends ComBat to unseen sites using a prior fit, avoiding data sharing similar to CREB. This was primarily demonstrated on cortical thickness rather than high dimensional FC, and did not demonstrate biological signal preservation (though that would likely be expected). This tool was developed in R and we have simultaneously but independently developed a tool in python. It is important to note that ComBat-Predict is a preprint, at the time of the writing of this manuscript, and this parallel development underscores the methodological relevance and timeliness of this approach within the field.

In this work, we introduce and validate CREB, a two-stage harmonization method. First, using CREB Learn, we generated a lightweight, distributable reference ‘bundle’ from large-scale, publicly available lifespan datasets, including the HCP Young Adult, Dallas LifeSpan Brain Study, and Southwest University Adult Lifespan Dataset. We then evaluated CREB Apply by comparing its output to the established NeuroHarmonize method. Our results show that CREB’s output is highly similar to NeuroHarmonize in both euclidean distance and mean absolute error. Furthermore, one-way ANOVA confirms that CREB effectively removes site-related variance that performed on par with NeuroHarmonize, while successfully preserving biological associations between neuroimaging features and age. We primarily computed harmonization on connectivity data but we then also showed that CREB was also able to harmonize total gray matter volume. We therefore present CREB as an effective harmonization tool whose distributable reference point makes it uniquely suited for machine learning workflows by preventing data leakage and enabling the seamless harmonization of new, unseen data. The CREB software is publicly available on [Github](https://github.com/tetra-tools/CREB).

## 2 Methods

### 2.1 MRI data

We downloaded open-source or publicly available datasets that had a structural T1-weighted MRI and resting-state fMRI data; we downloaded data from “healthy control” participants from each set. Below we describe a list of studies that we preprocessed and used for data analysis in this manuscript. Information on each study and MR sequence information is presented in table 1 - the train and test data are split. For each dataset, we additionally have a document that describes in greater detail each dataset, their inclusion/exclusion criteria, and MR acquisition - these are located in supplemental documents 1-12. All data was downloaded, converted to Brain Imaging Data Structure (BIDS) format, and then preprocessed.

**Table 1:** Participant’s demographic and MRI parameters of each dataset used in the analysis. TR: Repetition Time; TE: Echo Time. The TR column in Table 1 for CamCAN dataset contains both values for resting task and movie task. The Rest, Non-Rest column contain runs that we used as input for our pre-processing. Since Aging is multi-echo dataset, the TE columns contains all 3 echo times.

| **Train Dataset** | **Age (yrs)** | **%Female** | **TR (ms)** | **TE (ms)** | **Fieldmap** | **# Rest/# Non-Rest** |
| --- | --- | --- | --- | --- | --- | --- |
| HCP-YA | 22-37 | 54.4% | 720 | 33.1 | Yes | 1083, 0 |
| BVDCC | 20-86 | 62.7% | 2000 | 27.0 | No | 157, 470 |
| DLBS | 21-97 | 61.9% | 2000 | 30.0 | No | 1442, 5025 |
| SALD | 19-80 | 62.2% | 2000 | 30.0 | No | 792, 0 |
| Bilingual | 18-52 | 76.6% | 1500 | 30.0 | Yes | 64, 0 |
| UCLA-CNP | 21-50 | 43.0% | 2000 | 30.0 | No | 1298, 0 |
| Music | 21-70 | 100.0% | 3000 | 40.0 | No | 156, 0 |
| Placebo | 44-78 | 52.6% | 2500 | 30.0 | No | 76, 0 |
| Depression | 19-55 | 73.6% | 2500 | 35.0 | No | 62, 0 |
| **Testing Dataset** | **Age (yrs)** | **%Female** | **TR (ms)** | **TE (ms)** | **Fieldmap** | **# Rest/# Non-Rest** |
| CamCAN Rest+Movie | 18-88 | 50.5% | 1960 or 2470 | 30.0 | No | 605, 527 |
| Aging | 18-83 | 56.1% | 3000 | 13.7, 30.0, 47.0 | No | 592, 0 |
| Glia | 30-84 | 57.9% | 3000 | 30.0 | No | 188, 0 |

#### 2.1.1 Training Data

[HCP Young Adult](https://db.humanconnectome.org/) (HCP-YA). This study focused on mapping the healthy human connectome by collecting and freely distributing neuroimaging and behavioral data on 1,200 normal young adults, aged 22-35 years. We used the 1200 Subject release version of this set. We downloaded data from 1206 participants - each participant had just one visit. We downloaded T1-weighted structural, T2-weighted structural, resting state fMRI, and fieldmaps. We used just the first session’s first run from each participant. All scans were collected on a 3T Siemens Skyra Connectom scanner at the Washington University in St. Louis. Greater detail is described in supplemental document 1.

[BOLD variability during cognitive control for an adult lifespan sample](https://openneuro.org/datasets/ds005270/versions/1.0.0) (BVDCC). The purpose of this study was to examine age-related changes in large-scale brain networks and their reduced specificity during cognitive control tasks across different domains in older adults. We downloaded data from 158 participants each with a single MR session. We downloaded T1-weighted, T2-weighted Fluid-Attenuated Inversion Recovery (FLAIR), and both resting state and non-resting state fMRI. Tasks included were the n-back Working Memory, Go/No-go, and Transportation switch. MR data were collected using Siemens Trio 3T scanner. Greater detail is described in supplemental document 2.

[Dallas Life Brain Study](https://fcon_1000.projects.nitrc.org/indi/retro/dlbs.html) (DLBS). This study was designed to integrate brain and cognition across the adult lifespan. We downloaded data from 464 participants - each participant had up to 3 follow-up scans with 4-5 years in between scans. We downloaded T1-weighted, T2-weighted Fluid-Attenuated Inversion Recovery (FLAIR), and both resting state and non-resting state fMRI. The study had several fMRI tasks including the Ventral Visual Task, Words Task, Scene Task, and resting state. Some tasks have multiple runs, and we used all runs. All participants in the first and second timepoints were scanned on a single 3T Philips Achieva Scanner at the University of Texas Southwestern Medical Center (UTSW) while the third timepoint was completed on a newer 3T Achieva Scanner at UTSW. Greater detail is described in supplemental document 3.

[Southwest University Adult Lifespan Dataset](https://fcon_1000.projects.nitrc.org/indi/retro/sald.html) (SALD). This study was designed to study the developmental trajectory of the human brain and changes in network function in aging. The dataset contains 494 participants each with a single MR session. We downloaded T1-weighted structural resting state fMRI for each participant. All data was collected at the Southwest University Center for Brain Imaging using a 3T Siemens Trio MRI scanner. Greater detail is described in supplemental document 4.

[Bilingualism and the brain](https://openneuro.org/datasets/ds001796/versions/1.7.0) (Bilingual). This study was designed to examining neurological effects of different individual language differences in bilingual adults. We downloaded data from 64 healthy participants each with a single MR session. We downloaded T1-weighted structural images, resting and non-rest fMRI, and fieldmaps. The only task was the flanker task and we did not include the non-rest runs. MR images were collected on a 3T Siemens Prisma scanner. Greater detail is described in supplemental document 5.

[UCLA Consortium for Neuropsychiatric Phenomics LA5c Study](https://openneuro.org/datasets/ds000030/versions/00016) (UCLA-CNP). This study focused on understanding memory and cognitive control (response inhibition) function in both healthy individuals (130 subjects) and individuals with neuropsychiatric disorders including schizophrenia (50 participants), bipolar disorder (49 participants), and attention deficit/hyperactivity disorder (43 participants). We downloaded data from 272 participants who each have a single MR scan. We only used data from healthy controls. We used T1-weighted structural images and fMRI from each participant that included resting and non-resting state fMRI. Non-rest tasks included were: Balloon Analog Risk Task (bart), Breath-Holding Task (bht), Paired Associates Memory Task - Encoding (pamenc), Paired Associates Memory Task - Retrieval (pamret), Spatial Working Memory Capacity Tasks (scap), Stop-Signal Task (stopsignal), Task Switching (taskswitch). MR images were collected on a 3T Siemens Trio scanner. Greater detail is described in supplemental document 6.

[Functional Connectivity of Music-Induced Analgesia in Fibromyalgia](https://openneuro.org/datasets/ds001928/versions/1.1.0) (Music). This study was designed to understand the analgesic properties of music on chronic pain. We downloaded data from 20 participants with Fibromyalgia and 20 healthy controls, but we only used data from healthy controls. Each participant had 4 runs of resting fMRI before and after music and before and after pink noise. MR data were collected with GE Discovery 750W 3T MRI Scanner. Greater detail is described in supplemental document 7.

[Brain connectivity predicts placebo response across chronic pain clinical trials](https://openneuro.org/datasets/ds000208/versions/1.0.1) (Placebo). This is a cross-sectional dataset designed to study if clinical placebo response can be predicted from resting state connectivity. There were total of 76 participants with 20 healthy controls; we only use data from controls. Participants all had a T1-weighted structural scan and a resting state fMRI. All MR data were collected using a 3T Siemens Trio in the Northwestern University, Feinberg School of Medicine. Greater detail is described in supplemental document 8.

[Resting state with closed eyes for patients with depression and healthy participants](https://openneuro.org/datasets/ds002748/versions/1.0.5) (Depression). This is a cross sectional dataset designed to study resting state network differences between depressed and healthy participants. There were a total of 72 participants in this study with 21 healthy controls - we only used data from controls. Participants had a T1-weighted structural scan and resting state fMRI with eyes-closed. All MR data were collected using 3T Philips Ingenia scanner in International Tomography Center, Novosibirsk. Greater detail is described in supplemental document 9.

#### 2.1.2 Testing Data

[Cambridge Center of Aging and Neuroscience](https://cam-can.mrc-cbu.cam.ac.uk/dataset/) (CamCAN). We used CamCAN, a cross sectional adult life span dataset in our analysis,. The main goal of this study was to characterize age-related changes in cognition, brain structures and functions to study the neurocognitive mechanism behind healthy brain aging. There were a total of 653 participants. We downloaded T1-weighted and T2-weighted structural, resting state fMRI, and movie watching fMRI. All scans were collected using a 3 T Siemens TIM Trio scanner with a 32-channel head coil in the Medical Research Council (UK) Cognition and Brain Sciences Unit (MRC-CBSU) in Cambridge, UK. Greater detail is described in supplemental document 10.

[Neurocognitive aging data release with behavioral, structural, and multi-echo functional MRI measures](https://openneuro.org/datasets/ds003592/versions/1.0.13)(Aging). This dataset was designed to study neurocognitive aging. There were a total of 301 participants from 2 sites. For resting state connectivity harmonization, we only used data from 1 site - which resulted in 231 participants; for structural data harmonization, we used both sites. We downloaded T1-weighted, T2-weighted Fluid-Attenuated Inversion Recovery (FLAIR), and resting state fMRI. Each participant had two resting state scans and one of which was a multi-echo fMRI sequence. Site 1 had participants scanned with a 3T GE Discovery MR750 and 32-channel head coil at the Cornell Magnetic Resonance Imaging Facility, and site 2 had participants scanned with 3T Siemens Tim Trio MRI scanner with a 32-channel head coil at the York University Neuroimaging Center in Toronto. Greater detail is described in supplemental document 11.

[A large-scale dataset of pre- and post-surgical MRI data in patients with chronic trigeminal neuralgia](https://openneuro.org/datasets/ds005713/versions/1.0.1) (Glia). This dataset was design to study resting state connectivity changes after surgery of patients diagnosed with chronic trigeminal neuralgia. We downloaded data from 112 patients with trigeminal neuralgia and 48 healthy controls - we only used data from controls. The patient group had two sessions but the controls only had a single MR session. We downloaded T1-weighted structural data and resting state fMRI. The MR data were collected using 3T Philips Ingenia scanners in the Federal Neurosurgical Center in Novosibirsk. Greater detail is described in supplemental document 12.

### 2.2 MRI Preprocessing

#### 2.2.1 Functional Images

The MRI preprocessing was done with a singularity image of [fmriprep-23.2.3](https://fmriprep.org/en/23.2.3/) and singularity image of [xcpd-0.10.5](https://xcp-d.readthedocs.io/en/0.10.5/). For fmriprep, we disabled all freesurfur related processing and ran fieldmap-less susceptibility-derived distortions estimation for all datasets that did not contain a fieldmap. If a dataset had skull-stripped T1-weighted images (e.g. Depression Dataset), we skipped skull stripping. The fmriprep output space was MNI152NLin2009cAsym. The processing details were further detailed in the supplemental document 13 (fmri prep) and 14 (xcpd); we briefly describe the processing here.

Briefly, fmriprep conducted the following preprocessing: T1-weighted bias correction using N4BiasFieldCorrection [Tustison et al., 2010]. This was skull-stripped using antsBrianExtraction. Brain tissue segmentation of cerebrospinal fluid (CSF), white-matter (WM) and gray-matter (GM) was performed on the brain-extracted T1w using fast (in FSL, [Zhang et al., 2002]). Spatial normalization was conducted using nonlinear registration with antsRegistration.

For functional processing, this was done similarly across studies and sequences (including for task). A high contrast reference image was first generated and head-motion parameters with respect to this reference (transformation matrices, and six corresponding rotation and translation parameters) were estimated before any spatiotemporal filtering using mcflirt [Jenkinson et al., 2002]. Slice timing correction was applied using 3dTshift in AFNI - this was only done for Depression, Aging, UCLA-CNP, Music, Bilingual, SALD, BVDCC, CamCAN datasets in which slice timing was present in the dataset’s JSON side card, but was not done for Placebo, Glia, DLBS, HCP-YA, datasets. We conducted susceptibility distortion correction - this generated a phase difference map that can be used for susceptibility correction. For the studies that do not have fieldmaps, we used fieldmap-less correction. The fMRI reference image was then registered to the T1 image using mri coreg in freesurfer and boundary-based registration in FSL [Jenkinson and Smith, 2001]. We computed the framewise displacement (FD) using the Power et al estimation [Power et al., 2014]. We extracted signals from the CSF and WM using principal components analysis using anatomical comp corr (aCompCor) - which removes a probable GM mask. Components were calculated separately within the WM and CSF masks. This was extracted after conducting a high-pass filter with 1/128 Hz (0.008). For each CompCor decomposition, the *k* components with the largest singular values were retained, such that the retained components’ time series were sufficient to explain 50 percent of variance across the nuisance mask. The head-motion estimates calculated in the correction step were placed within the corresponding confounds file. The confound time series derived from head motion estimates were expanded with the inclusion of temporal derivatives and quadratic terms for each [Satterthwaite et al., 2013]. Frames that exceeded a threshold of 0.5 mm FD were annotated as motion outliers. All resamplings were performed with a single interpolation step by composing all the pertinent transformations (i.e. head-motion transform matrices, susceptibility distortion correction when available, and co-registrations to anatomical and output spaces). Gridded (volumetric) resamplings were performed using nitransforms.

For XCP-D, we specified the input type as fmriprep and mode as none to set our custom parameters. We set the number of initial scans to be considered non-steady state as auto, using aCompCor as a nuisance regressor. We set 240 seconds as the minimun amount of time required after motion scrubbing to perform further post-processing. We set despiking to None, and conducted motion censoring with FD threshold set to 0.5mm. Each censored volume was regressed out. We set the full-width at half-maximum of the smoothing kernel to 8 and set a minimum coverage required for any parcel to 0. We set the lower band-pass filter to 0 and upper band-pass filter to 0.15 with a band-pass filter order set to 2. Note that we set the lower band-pass to 0 as we performed high-pass filtering in fMRIprep with 0.008Hz. For this analysis, we used timeseries from the Schaefer atlas supplemented with subcortical structures (4S) atlas [Schaefer et al., 2018, Pauli et al., 2018, King et al., 2019, Najdenovska et al., 2018, Glasser et al., 2013] for a total of 456 parcels. We computed the Pearson correlation matrix of each run by calculating all pairwise correlations.

#### 2.2.2 Anatomical Images

We used the output of [fmriprep-23.2.3](https://fmriprep.org/en/23.2.3/) to calculate gray matter volumes from the probability segmentation for each participant. For longitudinal studies with multiple session like DLBS, we added a bids filter so that fmriprep only processed the first session to make sure our resulting gray matter volume estimation was from a single session for each participant. We used a 0.5 threshold on the gray matter probability segmentation in native space. No augmentation of any kind was done on the gray matter volumes estimated and we only included participants who were healthy controls and had at least one included fMRI run that was not filtered out due to high motion for each dataset. Table 2 shows number of participants included, excluded, and percentage of healthy control participants we ended up using for each study. For studies designed for psychiatric disorders like UCLA-CNP, the T1w healthy control rate was low because a large portion of the participants were not healthy controls.

**Table 2:** Number of BOLD (both rest and non-rest) and structural images included/excluded and percentage included for each dataset. The structural data were derived with the fmriprep anatomical only pipeline. For T1w, we excluded participants who were not healthy controls or who do not have included fMRI runs.

| **Train Dataset** | **BOLD #(Inc/Exc)** | **BOLD Included %** | **T1w #(Inc/Exc)** | **T1w Healthy Control Included %** |
| --- | --- | --- | --- | --- |
| HCP-YA | 1053, 30 | 97.23% | 1037, 59 | 94.6% |
| BVDCC | 602, 25 | 96.01% | 152, 6 | 96.20% |
| SALD | 452, 34 | 93.00% | 454, 41 | 91.71% |
| DLBS | 6220, 247 | 96.18% | 258, 61 | 80.89% |
| UCLA-CNP | 1223, 167 | 86.3% | 112, 153 | 42.26% |
| Placebo | 59, 17 | 77.63% | 18, 48 | 27.27% |
| Bilingual | 64, 0 | 100% | 64, 0 | 100% |
| Music | 154, 2 | 98.72% | 20, 19 | 51.28% |
| Depression | 62, 0 | 100% | 20, 52 | 27.78% |
| **Test Dataset** | **BOLD #(Inc/Exc)** | **BOLD Included %** | **T1w #(Inc/Exc)** | **T1w Healthy Control Included %** |
| CamCAN | 1004, 128 | 88.70% | 491, 159 | 75.54% |
| Aging | 589, 3 | 99.49% | 295, 5 | 98.33% |
| Glia | 122, 66 | 64.89% | 29, 67 | 30.20% |

### 2.3 MRI Quality Control

We performed quality control on the motion FD outliers. For all runs, we evaluated and excluded by 3 criteria: (1) scans that had less than 3min of data after excluding outliers were not used; (2) scans that had more than 20% of volumes as outliers were not used; and (3) scans that contained any 1-min window with more than 50% outliers were excluded. This ensured that high quality data with low levels of motion were included. Table 2 describes the number of excluded runs for each study. It is importantly to note that the included and excluded runs were for rest and non-rest data combined; runs from non-healthy control were also counted if they pass the motion filtering criteria so the table gives an overview of image quality (in terms of motion) of the study. These runs from non-healthy control participants were excluded later. Some datasets contained additional quality metrics, and so we applied our motion exclusion in addition to this measure. For example, UCLA-CNP contained a column that indicated if images had ghosting artifacts, in such cases, we directly applied the provided quality metric and exclude all runs listed as having image quality issues.

### 2.4 Data Augmentation

We performed data augmentation on the fMRI runs by computing resting state connectivity with a continuous subset of the original timeseries. This augmentation is conducted 4 times on all runs to increase the size of the training data. For each run, we randomly selected a continuous chunk of the timeseries of variable starting point and duration. The minimum duration was always 3min for all dataset and the maximum duration depended on the total time of the particular run we were augmenting. These chunks of continuous timeseries from the original run were then used to generate a connectivity matrix, allowing for random variability in the connectivity matrix and augments the data. Thus, in addition to the connectivity matrix generated from the full run, participants had 4 more connectivity matrices generated that represented a matrix from a random chunk of their data with variable duration (at least 3 min). This generates ‘noisy’ connectivity data that can be used for augmentation. Additional details are included in the supplemental methods document 15.

### 2.5 Imputation

Due to poor coverage or low signal-to-noise ratio, some regions in the timeseries from the fMRI have missing data. Across all studies in the training set, we computed the average and variance of the connectivity matrix across participants excluding missing values. We used this mean and variance to estimate 3000 synthetic matrices by estimating values from that distribution. This generates 3000 synthetic connectivity matrices that can be used for imputing missing data randomly. For each missing value in each run, we then chose a random index (1 to 3000) and imputed missing values. We describe this further in supplemental methods document 16.

### 2.6 NeuroHarmonize

For comparison, we conducted traditional harmonization using a well-established approach (NeuroHarmonize or NeuroComBat). We combined all training and testing datasets and conducted harmonization across all studies while preserving age and sex biological variables. We used the opensource PyPI package for [NeuroHarmonize](https://github.com/rpomponio/neuroHarmonize). This was built on NeuroComBat [Pomponio et al., 2020, Fortin et al., 2018]. The major improvement it has compared to NeuroComBat was that it allows for fitting on non-linear covariates with spline. During the prior estimation and posterior update steps, NeuroHarmonize calls core functions from NeuroComBat to complete these procedures. This serves as a reference to show that CREB harmonization performs comparably to standard harmonization without data leakage.

### 2.7 CREB

Similar to NeuroComBat, we assume our raw, unharmonized data *y* is made up from baseline signal, biological signal, and site effects. We refer to all non-biological signal as ‘site’ effects though this can mean scanner, sequence, site, and other differences between datasets. We use *α*_*v*_ to denote baseline signal for each feature *v*, and 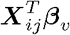 to denote the biological covariates for site *i* and participant *j*. ***X*** is an *n* ×*p* matrix of biological covariates, *β*_*v*_ is *p* ×1 vector of coefficient with ***X*** for feature *v, γ*_*iv*_ and *ϵ*_*iv*_ are the true additive and scaling site effects on site *i* on feature *v*, respectively. Thus, we have:

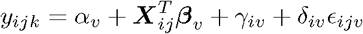

To harmonize the data, we estimate the site effect and remove them by subtracting the additive site effect and divide by the scaling site effect. Here we denote 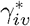 as our estimated additive site effect and 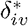 as the scaling site effect for site *i* and feature *v*.

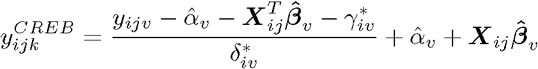

In CREB, our harmonization operates in two stages: CREB Learn conducts “bundle” generation and CREB Apply conducts harmonization. During CREB Learn, we take the training set and build a design matrix ***X*** containing only an intercept and biological covariates (age and sex). This ensures that site effects remain in the residuals rather than being regressed out prematurely. For each feature, we fit ordinary least squares and subtract these residuals across features and divide by the square root of pooled variance, producing standardized residuals that can be summarized at the site level. One critical difference between CREB and NeuroHarmonize is that our design matrix ***X*** only contains the intercept term and the covariate terms like age and sex. In NeuroHarmonize, the batches that each participant is coming from is one-hot encoded into the design matrix ***X*** in order to calculate the weighted-average baseline value for each feature. In CREB, this baseline *α*_*v*_ is handled with the intercept in the design matrix, hence we can write

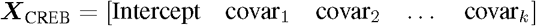

while in NeuroComBat/NeuroHarmonize, we have

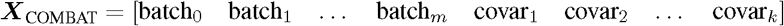

On the standardized data, for each site and feature, we compute sufficient statistics including sample mean 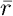, sample variance 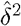, and sample size *n*), which are stored in the bundle along with the fitted regression coefficients, pooled variance, and feature metadata. From the bundle, we estimate global empirical Bayes priors. If we do joint update, we estimate the prior as a Normal-Inverse-Gamma distribution characterized by parameters (*κ*_0_, *a*_0_, *b*_0_); if we do iterative update like NeuroComBat, we estimate prior of *γ* as a Normal distribution with mean of *µ*_0_ and variance of 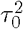, and estimate the prior of δ^2^ as an Inverse Gamma distribution with *α*_0_ and *β*_0_ using a method of moment estimator. These priors capture the mean and variance of the site effect across all training sites and features. It is important to note the distinction between CREB and NeuroHarmonize, in CREB the priors are estimated across both batch and features. Thus, in CREB, the training data generates a single set of priors and this is applied on all data we want to harmonize. In NeuroHarmonize, we would end up with one set of priors per site since this can be calculated directly as it has access to all of the data. When harmonizing new sites, we residualize and standardize data using the regression coefficients and pooled variance stored in the bundle, and then compute per-site residual’s summary statistics exactly as in the training. We assume after subtracting the biological signal, residuals are left with primarily site effect. For each site, we update the posteriors distribution of site effect using priors from the bundle. The mean of posterior distribution of site effect provides the additive site effect corrections and the mean of variance of posterior distribution of site effect provides the multiplicative site effect correction. This workflow is described in Figure 1.

**Figure 1.**
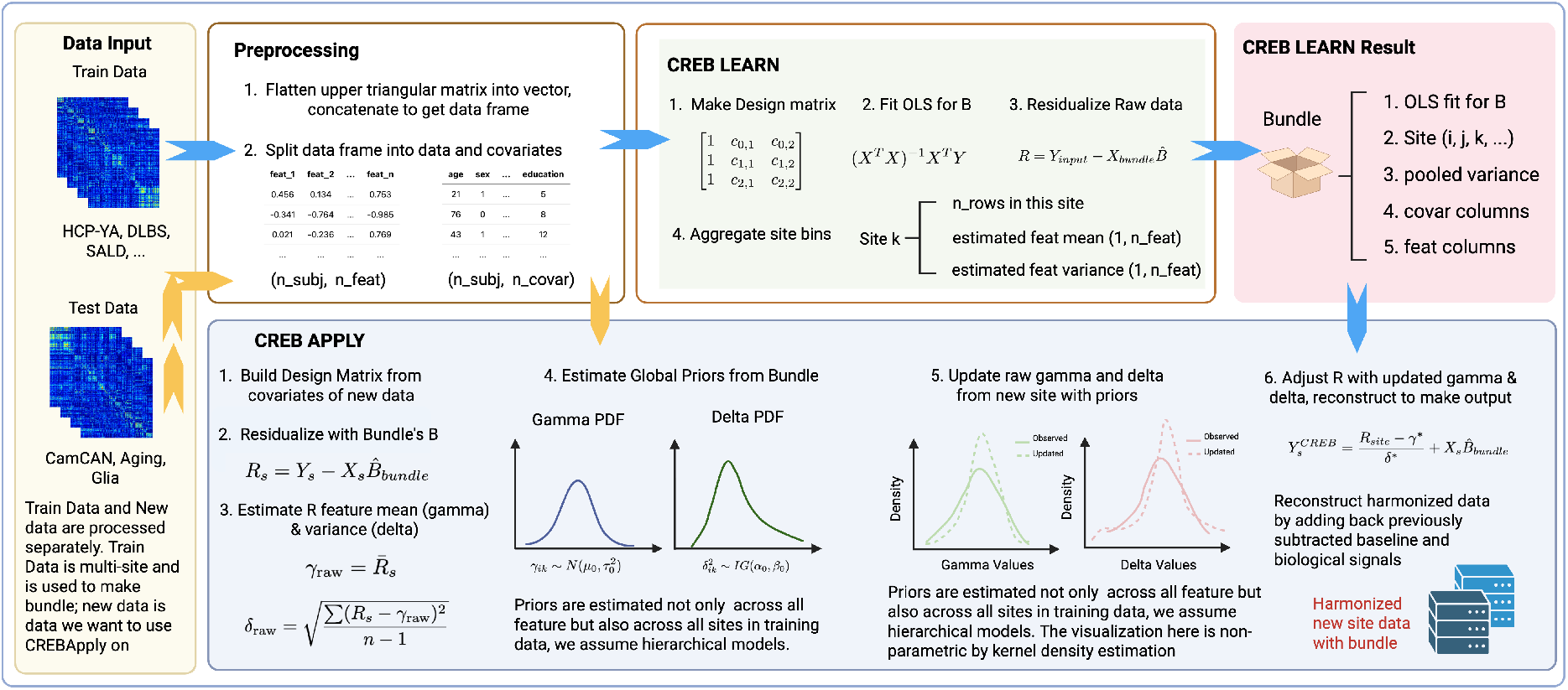
Workflow of CREB Harmonization. (Data Input) Train and test data are connectivity matrices across participants and fMRI sessions for task and rest. (Preprocessing) These matrices were flattened by extracting the upper triangular matrix. (CREB Learn) We fit a regression with biological signals (age and sex) and compute the residual on the raw connectivity data. We then compute the ‘bundle’ statistics on these residuals for estimating the prior distribution of site. (CREB Apply) In test data (e.g., CamCAN), we residualize the connectivity values using regression from train data. We then use the prior distribution and update the posterior for each new site we want to harmonize. We then reconstruct the harmonized data.

The pseudo-code for the closed form update is as listed in Algorithm 1. The joint update was done in one step where both mean and variance were estimated at the same time. This process ensured that site effects were shrunk towards the global prior, relative to the number of participants in a site, with small sites being shrunk more strongly, and large sites relying more heavily on their own data.

With updated posteriors, we remove the site effect by subtracting the site mean and dividing by site standard deviation. We then multiply back the pooled squared variance and add the biological signal to reconstruct harmonized data. In this algorithm, *n*_*s*_ is number of subjects in site we want to harmonize, 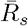 is the residual mean for each feature of this site, *κ*_0_, *α*_0_, and *β*_0_ are estimated priors from the training sites, *SST*_*s*_ is the sum of square total this site that we want to harmonize.

#### Algorithm 1

CREB Posterior Joint Update

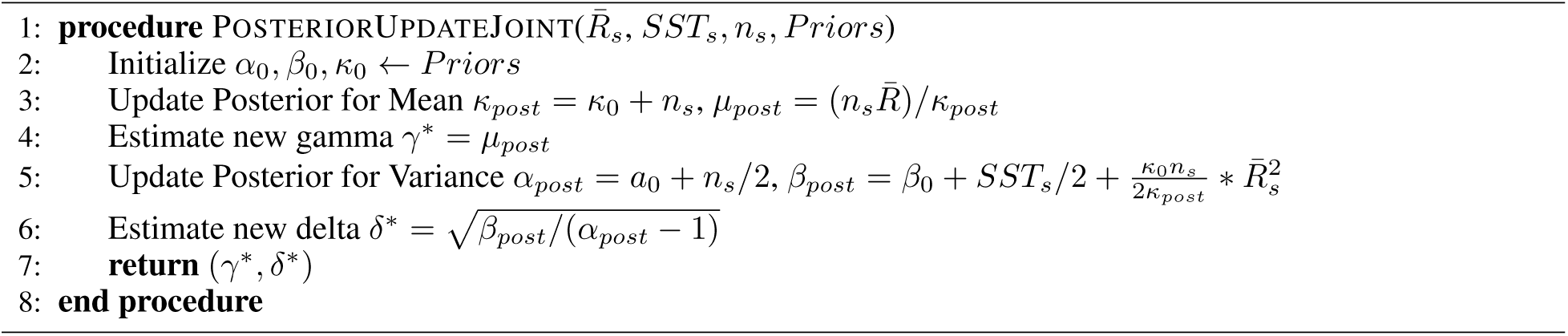

CREB can also do iterative updates like in the original ComBat [Johnson et al., 2007]. In iterative update, we have: 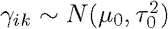 and 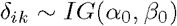. The priors *µ*_0_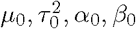, *α*_0_, *β*_0_ were calculated with methods of moment estimation. Importantly, unlike in NeuroCombat, we are not doing 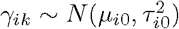 or 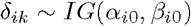 because our empirical Bayes estimation is not only leveraged across all features but also across all sites.

In the iterative update scheme, we estimate the new mean assuming that we know variance, and estimate variance with the newly updated mean. The iterative update process repeats until we reach a convergence point calculated from absolute difference between new and old mean and new and old variance. This is demonstrated in Algorithm 2. Like in joint update, the 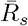 is residual feature mean of site *s* that we want to harmonize, *SST*_*s*_ is sum of square total of site *s, n*_*s*_ is size of site *s, R*_*s*_ is the residual of site *s*, we include it as a parameter input so we can calculate new *SST* value in the update process.

#### Algorithm 2

CREB Posterior Separate Update

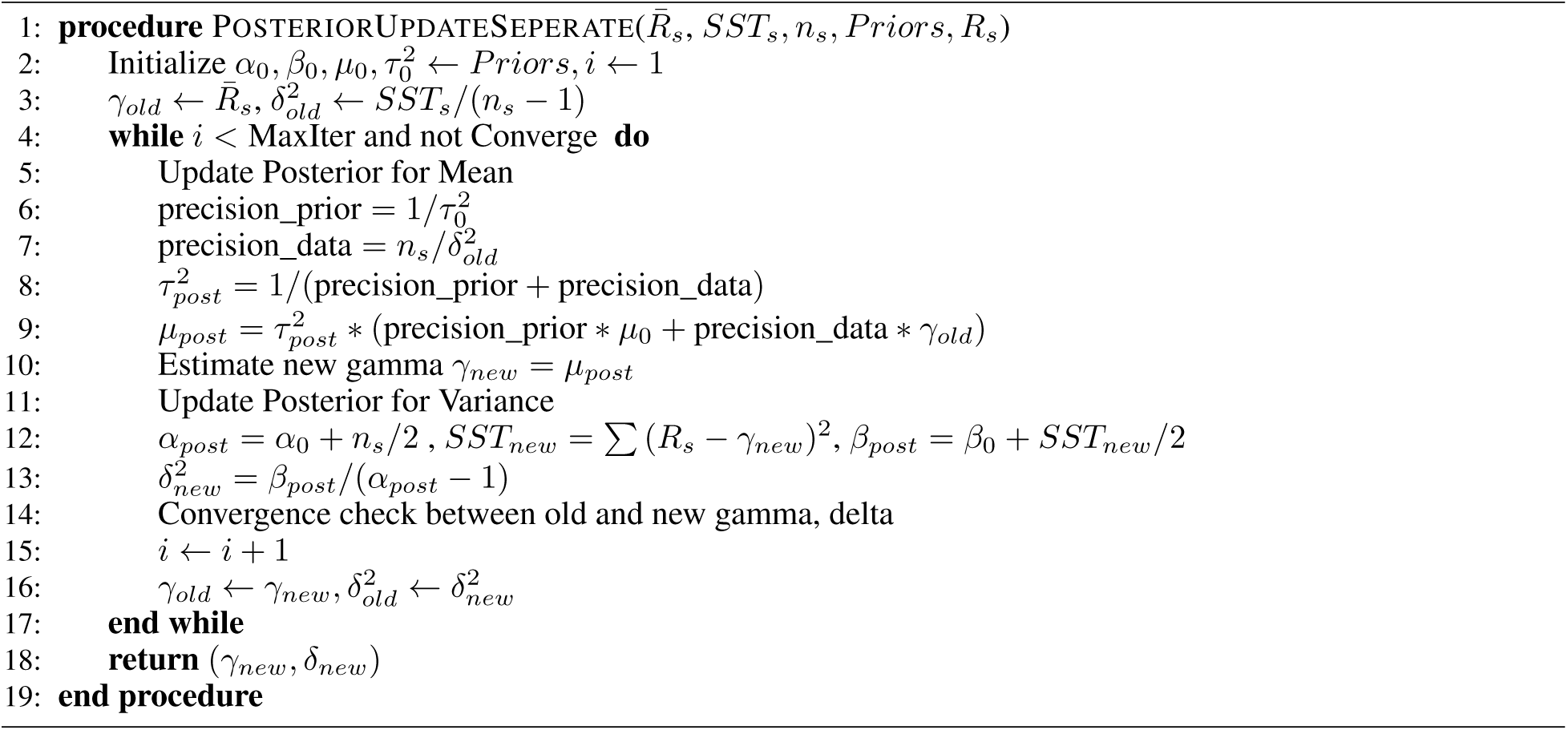

CREB Learn and CREB Apply allow for an easy two-stage workflow that first creates easily shareable bundles and then harmonizes new data. The harmonization step can be used with a bundle generated from other people as long as the data to be harmonized has the same covariates as data used to generate the bundle. The decoupling of prior with site in CREB allows for a single reference point for harmonization. In this way, completely new data can be easily harmonized with a deployed bundle and there is no re-harmonization needed with all previous sites every time a new site is included. This design is particularly useful for machine learning applications as it allows bundles to be generated once on a training set and safely applied to the validation, test, and external withheld testing cohorts without risk of data leakage. In addition, this bundle can be deployed easily along with the trained machine learning model so users who want to apply their data on the trained model can first harmonize their data. Users may select between the closed-form update, which estimates mean and variance simultaneously, or the iterative update, which alternates estimation of mean and variance until convergence. In our analyses, we used the closed-form update for connectivity features, and iterative update for gray matter volumes.

### 2.8 Statistical Analysis

We compared the harmonized outputs of CREB to that of NeuroHarmonize. CREB used the training data to estimate the priors and this was applied to the test set. NeuroHarmonize was run on train and test together - this is the standard approach to harmonization. The harmonization comparison is for both resting state connectivity from fMRI, and gray matter volume from T1w data.

On resting state connectivity, we compared site differences in the raw unharmonized data (RAW), CREB, and NeuroHarmonized data on a network level and edge-level using averaged Default Mode Network’s within-network connectivity (from XCPD’s 4S456 atlas which use Yeo 7Network[Yeo et al., 2011] for parcel’s network assignment) and medial prefrontal cortex (mPFC) to posterior cingulate cortex (PCC) connectivity since this is the canonical correlation for the DMN. For every edge in the test datasets, we computed the Euclidean distance and Mean Absolute Error (MAE) between each of the following pairs: (NeuroHarmonize, CREB) and (NeuroHarmonize, Raw) - this estimates the similarity in outcome between different harmonization approaches and put them in perspective with each other. To test removal of additive site effect and scaling site effect, we perform one-way ANOVA with site as the categorical variable and Bartlett’s Sphericity on a single edge of mPFC-x-PCC, DMN average connectivity, and all edges average connectivity for Raw, NeuroHarmonize, and CREB data from filtered testing sets which is not augmented, and with a single run per participant.

Successful harmonization must retain biological signals, so we tested this by fitting linear regression between connectivity with age for all edges on Raw, NeuroHarmonize, and CREB output from CamCAN data. We then calculated Pearson’s r value of all linear regression and compared the distribution of these r value between harmonization methods and with Raw.

We also conducted this on total gray matter volume as estimated from the T1-weighted image. We plotted the box-plot for each site for Raw, NeuroHarmonize, and CREB for training sites, and external test sites. We tested biological preservation by fitting linear regression between the gray matter volume and age and calculate the *r*^2^ value for each harmonization method.

## 3 Results

### 3.1 Harmonization Similarity

Figure 2 shows the mPFC-x-PCC connectivity for the training set that we used in bundle generation (Figure 2A) and for testing set (Figure 2B) of raw data as well as harmonized outputs from NeuroHarmonize and CREB. Figure 2C and 2D show the average DMN connectivity for train and test sets, respectively. These box-plots show that both NeuroHarmonize and CREB perform similarly well, each harmonizing to their own “mean.” Both approaches harmonize between sites. CREB was able to harmonize the test studies CamCAN, Aging, and Glia in a two-stage process that prevents leakage.

**Figure 2.**
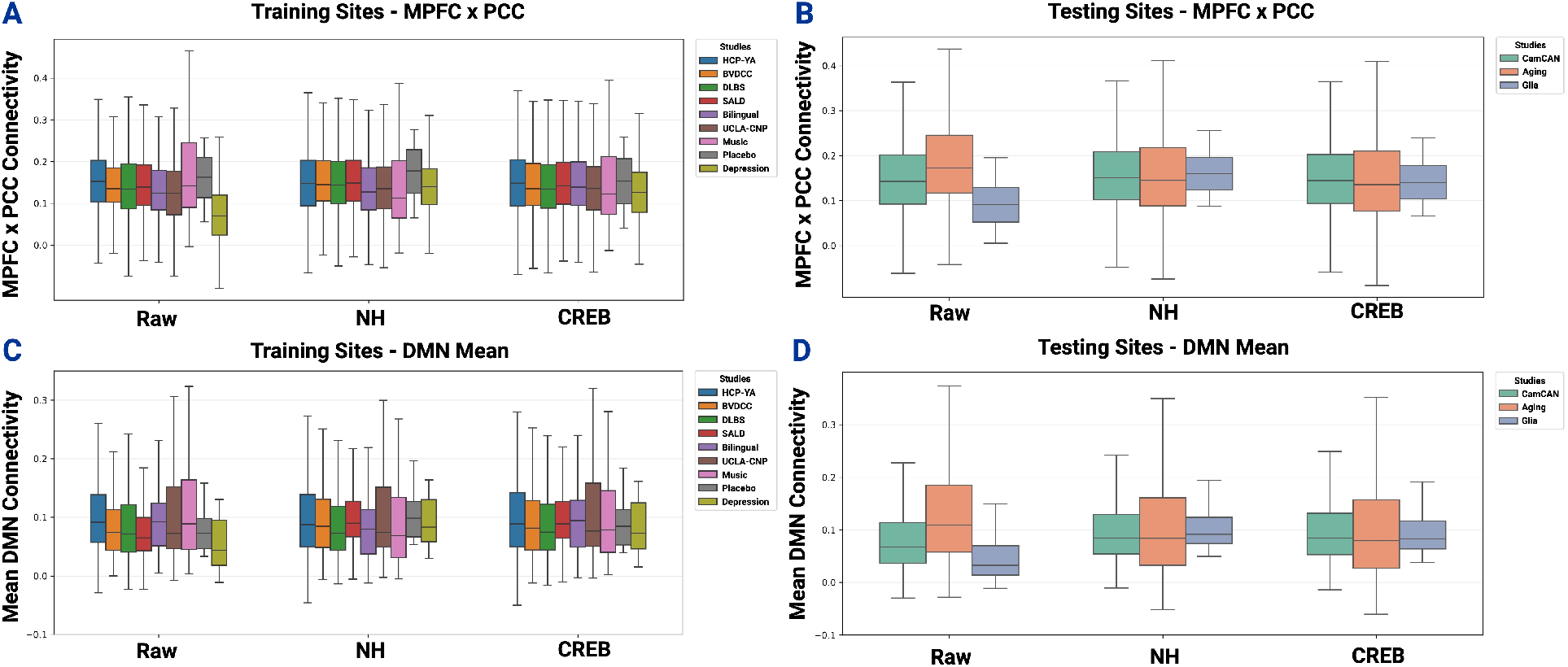
Box-Plots showing connectivity of the Raw data, NeuroHarmonize (NH), and CREB for mPFC-x-PCC connectivity for train (2A) and test (2B) as well as DMN connectivity for train (2C) and test (2D).

In Figure 3, we highlight differences in NeuroHarmonize and CREB (Figure 3A), where NeuroHarmonize conducts harmonization on train and test datasets together while CREB does this in two steps (learn and apply) that prevent leakage. The average Euclidean distance between NeuroHarmonize and CREB across all edges was 2.6, the distribution is shown in Figure 3B. The average MAE between NeuroHarmonize and CREB was 0.019 - this distribution is shown in figure 3C and highlights a max difference of approximately 0.08. This figure establishes that CREB produces output similar to NeuroHarmonize output, which is the mainstream method for harmonization.

**Figure 3.**
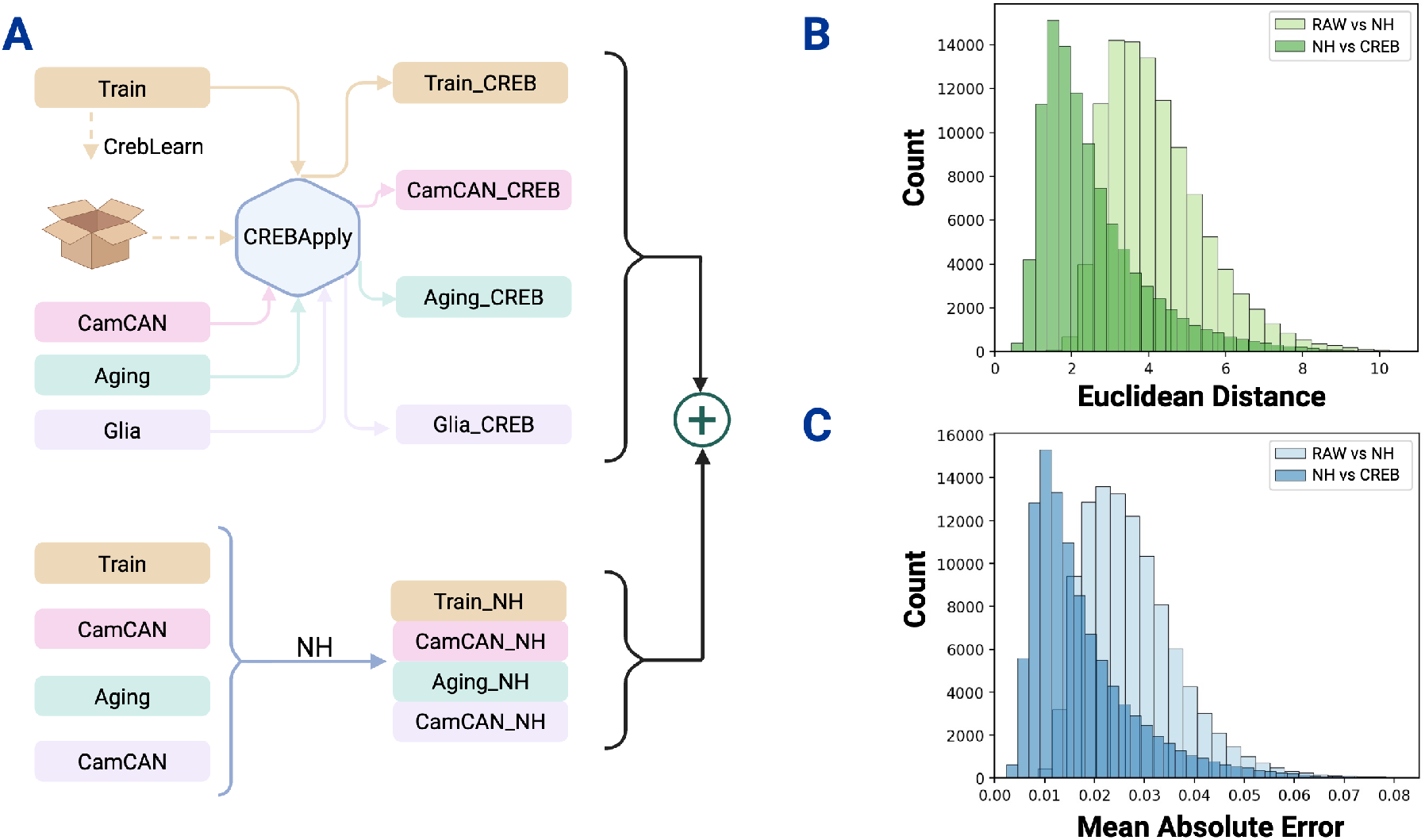
Comparison between NeuroHarmonize and CREB. The differences in methods are shown (3A) to highlight the two-step nature of CREB. (3B) We compute the Euclidean distance between Raw and NeuroHarmonize (NH) as well as NH and CREB. (3C) Mean absolute error between Raw and NH as well as NH and CREB. These plots show that both NH and CREB were able to harmonize data successfully.

### 3.2 Site Separability

We computed a one-way ANOVA on the mPFC-x-PCC connectivity, average DMN connectivity, and the global average connectivity for test datasets. We found that the site effect was significant for all three measures on Raw data (*F* (2, 813) = 26.3, *p* < 0.001; *F* (2, 813) = 45.5, *p* < 0.001; and *F* (2, 813) = 47.2, *p* < 0.001, respectively). However, there were no site differences in either NeuroHarmonize (*F* (2, 813) = 0.04, *p* = 0.965; *F* (2, 813) = 0.8, *p* = 0.472; *F* (2, 813) = 0.6, *p* = 0.571, respectively) or CREB (*F* (2, 813) = 0.8, *p* = 0.431; *F* (2, 813) = 1.1, *p* = 0.321; and *F* (2, 813) = 1.3, *p* = 0.283, respectively). We then computed a one-way ANOVA on each feature separately for all three methods for additive site effects, and then conducted a false-discovery rate (FDR, Benjamini-Hochberg) correction for multiple comparisons. In the raw data, 89643 edges (out of approximately 103*k* edges) were significantly different between sites. We conducted the same test in NeuroHarmonize data and found that only 3 edges were significantly different between sites. On CREB harmonized data, we found 0 edges were significantly different between sites. The result shows site effect removal for both NeuroHarmonize and CREB. We tested for scaling site effects with a Bartlett’s Sphericity test on every feature and corrected with FDR. In the Raw data, we found that 40566 edges were significantly different between sites. In NeuroHarmonized and CREB harmonized data, we found 11 and 0 edges that showed significant site differences, respectively.

### 3.3 Preserving Biological Signal

To investigate whether the biological signal could be preserved for functional connectivity, we conducted the following analysis on CamCAN alone. We computed the top-edge regression and found that the ROI pair of Left Hemisphere Visual Network 6 x Left Hemisphere Visual Network 19, and Left Hemisphere SomatoMotor Network 5 x Left Hemisphere SomatoMotor 35 had the strongest correlations between connectivity and age with the raw *r*^2^ values being 0.18, and 0.17, respectively. After NeuroHarmonize, the biological signal was largely preserved with the *r*^2^ values being 0.18, and 0.16, respectively. After CREB harmonization, the biological signal was largely preserved, with the *r*^2^ values for the three edges being 0.18, and 0.13, respectively - these are shown in in Figure 4

**Figure 4.**
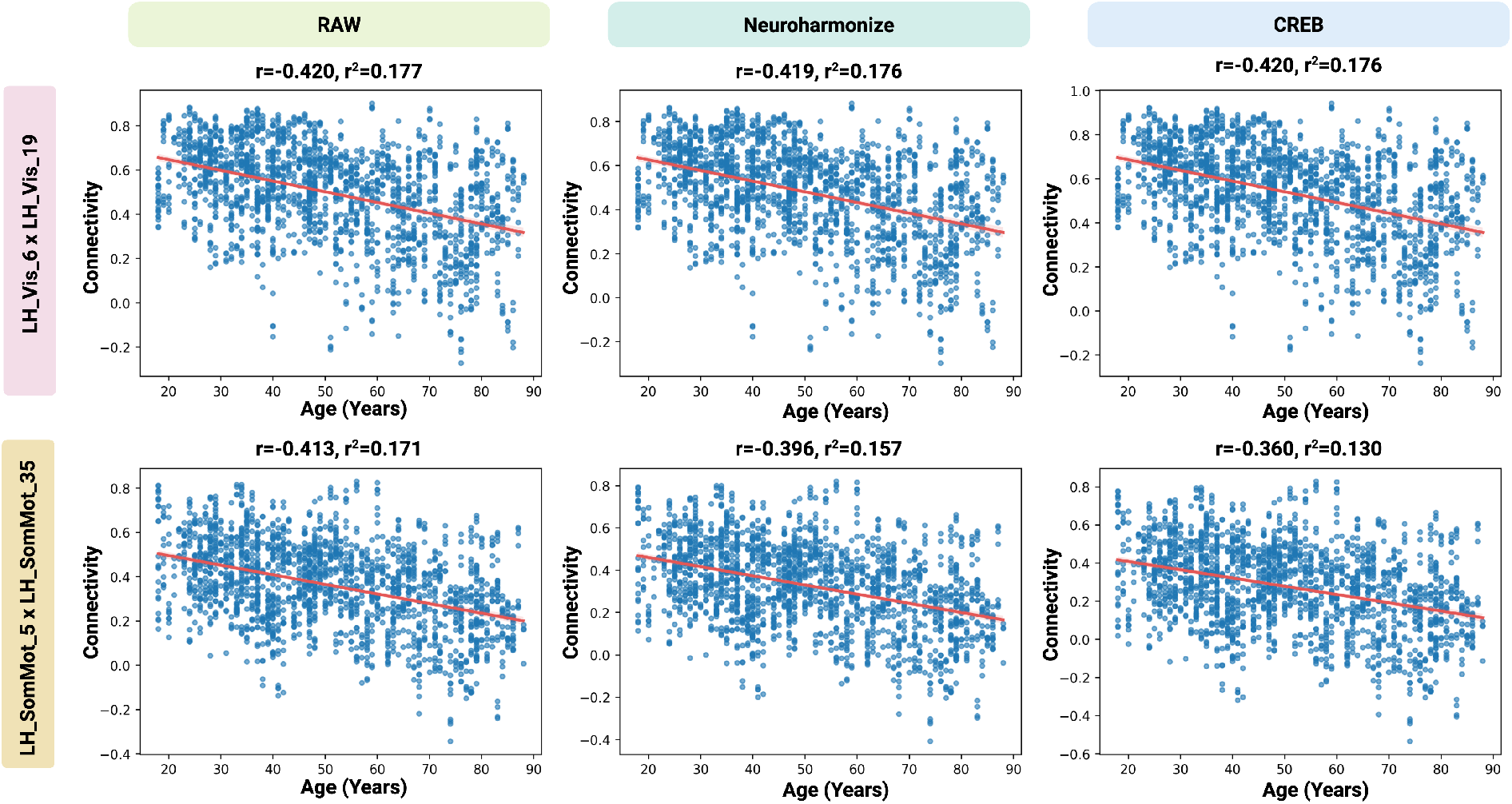
Biological signal preservation from linear regression of single edge with age. We show that the biological signal is preserved from raw data to NeuroHarmonize and CREB for two different edges.

We then repeated this analysis for every edge. We generated a difference density plot and violin plot of the distribution of Pearson’s r value across all edges in Figure 5. The plots show that these r values were highly preserved in each method of harmonization (i.e., there was no change in association from Raw to post-harmonization).

**Figure 5.**
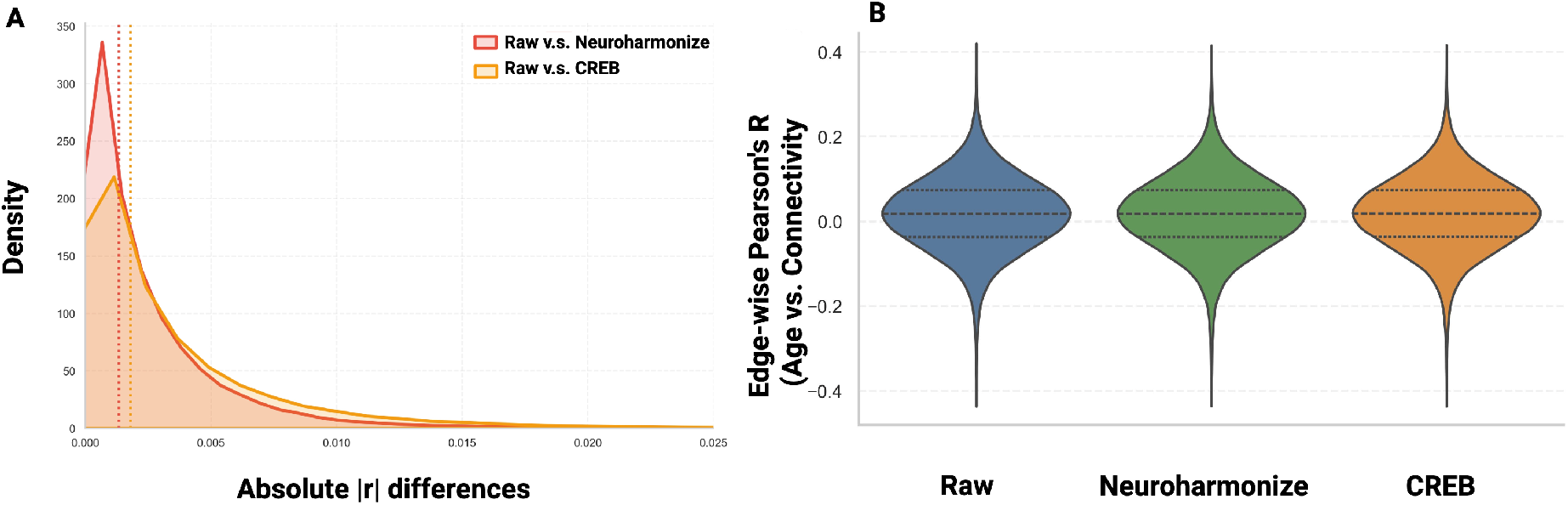
We conducted a regression with age across all edges. (A) We show absolute R differences for Raw and Neuroharmonize and for Raw and CREB. This is the difference in absolute r for each feature. (B) We show a Violin plot of edge-wise Pearson’s r between raw, NeuroHarmonize, and CREB.

We also tested the preservation of biological signal using gray matter volume as the feature to harmonize. We calculated box-plots for both train and test data. It is important to note that since we used both sites from the aging dataset for the analysis on structural data, we have 4 external sites in Figure 6. We show that CREB and NeuroHarmonize were able to harmonize the data equally well.

**Figure 6.**
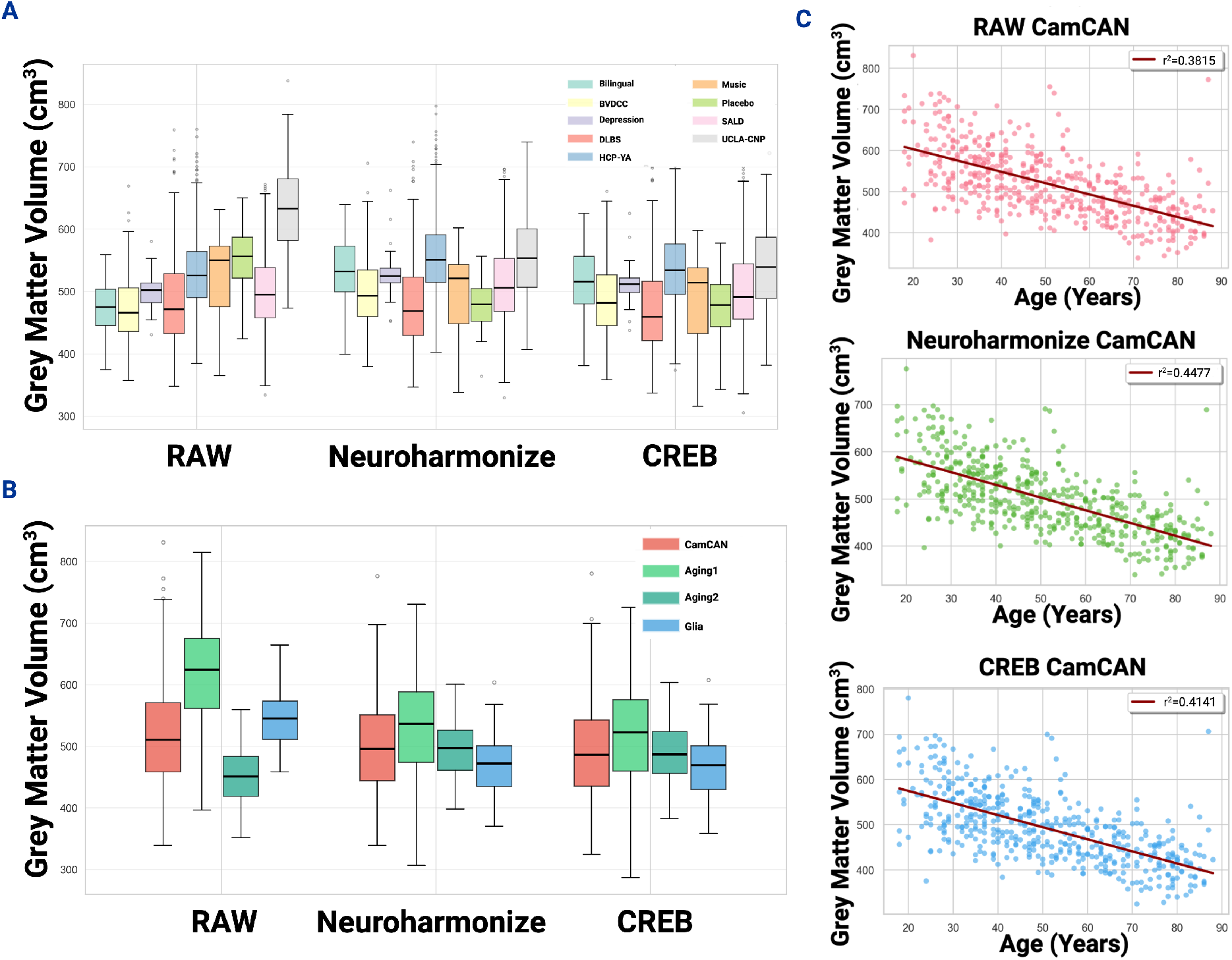
Harmonization of total gray matter volume across sites and biological signal preservation with age. We show box-plot of harmonization result between Raw, NeuroHarmonize, and CREB on train (6A) and test (6B) data. (6C) We show a scatter plot with fitted linear regression between age and total gray matter volume for different harmonization methods on the CamCAN dataset.

We fit a linear regression between age and gray matter volume on Raw, NeuroHarmonize, and CREB data. The *r*^2^ on Raw was 0.38, on NeuroHarmonize was 0.45, and on CREB was 0.41. This result shows that CREB was a generalizable method that can harmonize both functional connectivity and structural features; the result also further verifies CREB’s effectiveness in biological signal preservation. Additionally, we found that CREB performs similarly, in comparison to NeuroHarmonize in terms of site separability reduction. There was a significant reduction in site effect for both CREB (*F* (3, 811) = 7.1, *p* < 0.001, Bartlett’s *χ*^2^ = 26.2, *p* < 0.001 and NeuroHarmonize (*F* (3, 811) = 10.4, *p* < 0.001, Bartlett’s *χ*^2^ = 29.0, *p* < 0.001) when compared to Raw (*F* (3, 811) = 109.5, *p* < 0.001, Bartlett’s *χ*^2^ = 35.0, *p* < 0.001).

## 4 Discussion

We developed a new, two-stage harmonization method that enables harmonization of new, previously unseen sites to a single reference point established from training data. This reference point can be easily shared within a machine learning algorithm without sharing all of the training data - delivered as a ‘bundle.’ We found that CREB was able to harmonize unseen test data comparably to NeuroHarmonize, which had access to both train and test data. This approach avoids data leakage between train and test sets, but allows for harmonization of unseen test data. We showed that there were no site differences after harmonization with NeuroHarmonize or CREB, though NeuroHarmonize exhibited small site-specific differences on a few edges. Finally, we found that associations with biological signals (in this case, age) were preserved after harmonization with either approach. Thus, CREB is as effective as NeuroHarmonize at removing site effects in functional connectivity and gray matter volume data. CREB’s two-stage procedure allows for harmonization on the training data, which can then be applied to test data even those not available to the researchers. In addition, this can be deployed in a lightweight model as the “bundle” statistics take up 12.51MB. This approach addresses the issue of train-test data leakage during harmonizing.

Increased access to imaging due to data sharing and big data has made it possible to train machine learning models on vast datasets. One major challenge is that imaging data is highly impacted by ‘site’ effects including scanners, sequences, and field strengths, etc. There is compelling evidence that harmonizing data prior to training models improves their performance [Tassi et al., 2024]. One major challenge here is that harmonizing train and test data jointly can lead to data leakage [Rosenblatt et al., 2024] and potentially inflate performance of these models. Thus, there is a need to conduct this harmonization without leakage. We have shown that CREB is capable of conducting harmonization in a two-step procedure that harmonizes train and test sets independently. This can be deployed in machine learning models as a small ‘bundle’ (<13MB) to easily harmonize unseen test data.

Despite its strengths, our analysis has several limitations. During preprocessing, the DLBS (TR=2s), Glia (TR=3s), Placebo (TR=2.5s), and HCP-YA (TR=0.72s) datasets did not have slice time in their JSON side card. Given that HCP-YA’s TR is very short, we can probably safely preprocess it with fmriprep without doing slice timing correction, but for the other 3 dataset, doing fmriprep without slicetiming correction may be problematic. Of note, both Placebo and Glia dataset authors sent clarifications, but they arrived after we completed preprocessing all the data. One limitation of CREB is that a new bundle is needed for each fmriprep, xcpd, atlas choice and biological co-variate included. This is true of other harmonization approaches as well. While producing a new bundle for each preprocessing pipeline is expected, requiring new bundles for every user-defined combination of biological covariates is still a major limitation of most harmonization methods. Additionally, like other ComBat-style harmonization approaches, CREB assumes that the biological data distributions represented in the training data sufficiently overlap with those in the target dataset. This limitation is not unique to CREB and is inherent to harmonization methods that separate biological and site effects, and it becomes more pronounced for complex or highly structured covariates, where full covariate support across datasets is rarely available. Thus, it is recommended that the covariates in the train and test set have similar distributions. This is an assumption true of all harmonization approaches and is an important consideration here as well. This is especially important for biological variables that are quite complex and difficult to capture. For example, if we want to harmonize data while preserving depression biological signal, the training data should have a similar distribution on depression severity as the testing data. In machine learning, this would be expected since any biological variable that we train on needs to be well distributed, but nonetheless an important consideration to explicitly state. To further validate CREB’s performance in removing nuisance variance, future work could use traveling participants data to confirm that these methods maintain consistent subject-specific features despite variations in scanning parameters across sites.

Harmonization is critical for analyzing imaging data and for deploying generalizable machine learning models. CREB employs similar methods to ComBat to conduct data harmonization in a two-step procedure that prevents data leakage and is easy to deploy within machine learning models. CREB performed similarly to NeuroHarmonize both in terms of removing site specific effects but also in preserving biological variance. CREB is publicly available and can be easily integrated into machine learning workflows.

## Supporting information

supplement_01_hcp_ya

supplement_02_bvdcc

supplement_03_dlbs

supplement_04_sald

supplement_05_bilingual

supplement_06_ucla_cnp

supplement_07_music

supplement_08_placebo

supplement_09_depression

supplement_10_camcan

supplement_11_aging

supplement_12_glia

supplement_13_fmriprep

supplement_14_xcpd

supplement_15_creb_augmentation

supplement_16_creb_imputation

## 5 Acknowledgments

This project was not supported by any funding.

