## supplement_01_hcp_ya for "CREB: Consistent Reference External Batch Harmonization"

**Human Connectome Project Young Adult (HCP-YA) Detailed Description**

Study Design and Participants

The Human Connectome Project – Young Adult (HCP-YA)^1^ was a large study focused on characterizing brain connectivity and function in a population of 1200 healthy adults to enable detailed comparisons between brain circuits, behavior, and genetics at the level of individual participants. This was a cross-sectional study that recruited 1200 participants aged 22-35 years from the Missouri area. They recruited adult twins and non-twin siblings.

Inclusion criteria were age 22 to 35 years old and ability to give informed consent. Exclusion criteria were history of psychiatric disorder, substance abuse, neurological disorder, or cardiovascular disease; two or more seizures after age 5years or diagnosis of epilepsy; any genetic disorder; multiple sclerosis, cerebral palsy, brain tumor or stroke; head injuries; premature birth; currently on chemotherapy or immunomodulatory agents or history of radiation or chemotherapy; thyroid hormone treatment in the last month; treatment for diabetes except gestational or diet-controlled diabetes; migraine medication daily use; Folstein Mini Mental State Exam ≤ 25; moderate claustrophobia; pregnancy; or unsafe metal in body. The study was approved by local Institutional Review Boards. All participants gave written informed consent prior to participating.

MRI Acquisition

All scanning was done at Washington University in St. Louis using a customized 3T Siemens Connectome Skyra scanner equipped with a 32-channel receive-only head coil and a ‘body’ transmission coil designed by Siemens specifically for the smaller space available using the special gradients of these scanners. A subset of participants underwent 7T scanning at the University of Minnesota, however this data was not used here. Scans were collected over two days: day 1 session 1 collected two T1-weighted and two T2-weighted structural scans; day 1 session 2 collected two 15-minute resting state fMRI and three task fMRI; day 2 session 1 collected diffusion weighted scans; and day 2 session 2 collected two additional resting state fMRI and four task fMRI. For this analysis, we used only the first structural and resting state fMRI.

- T1-weighted magnetization prepared rapid gradient echo (MPRAGE) were collected with repetition time (TR)=2400ms, echo time (TE)=2.14ms, flip angle (FA)=8deg, field of view (FOV)=224 x 224, and 0.7mm^3^ isotropic resolution.
- T2-weighted Sampling Perfection with Application optimized Contrasts using different flip angle Evolution (SPACE) were collected with TR=3200ms, TE=565ms, FA=not reported, FOV=224 x 224, and 0.7mm^3^ isotropic resolution.
- T2*-weighted blood oxygen level-dependent (BOLD) images using a gradient-echo echoplanar imaging sequence were collected with TR=720ms, TE=33.1ms, matrix size=208 x 180, voxel size=2.0mm^3^ isotropic, FA=52 degrees, and multiband acceleration of 8.
- Diffusion weighted data were acquired with a multi-shell diffusion sequence with b-values of 1000, 2000, and 3000 s/mm^2^ with 90 directions each with TR=5520ms, TE=89.5ms, matrix size=210 x 180, voxel size=1.25mm^3^ isotropic, and FA=78 degrees.
