## supplement_02_bvdcc for "CREB: Consistent Reference External Batch Harmonization"

**BOLD variability during cognitive control for an adult lifespan sample (BVDCC) Detailed Description**

Study Design and Participants

The purpose of the BVDCC study^1^ was to understand how age differences in brain function influence aging and cognitive control. This was a cross-sectional study where they recruited 158 participants aged 20 to 86 years old from the greater Toronto area.

Participants were screened to be healthy (i.e., free from major psychiatric or neurological conditions with no history of head trauma, cognitively normal with Mini Mental State Examination >26, right-handed, fluent English speakers, with normal or corrected-to-normal vision. Participants’ informed consent was obtained in accordance with protocol approved by the Research Ethics Board at Baycrest Health Sciences Centre.

MRI Acquisition

All scanning was done at Baycrest Health Sciences Centre on a 3T Siemens Trio with a 12-channel Siemens head coil. Participants underwent a 1.5 hours MRI that included 1) T2-weighted Fluid-Attenuated Inversion Recovery (FLAIR), (2) 10-minute gradient echo-planar imaging (EPI) sequence resting state, (3) T1-weighted magnetization-prepared rapid gradient echo (MPRAGE), (4) three BOLD fMRI tasks, (5) diffusion weighted imaging, and (6) if time permitted, arterial spin labeling.

- T1-weighted magnetization prepared rapid gradient echo (MPRAGE) were collected with repetition time (TR)=4000ms, echo time (TE)=2.63ms, flip angle (FA)=9deg, field of view (FOV)=192 x 256, and 1mm^3^ isotropic resolution.
- T2-weighted Fluid-Attenuated Inversion Recovery (FLAIR) were collected with TR=3200ms, TE=466ms, FA=120deg, and 0.5x0.5x1mm^3^ resolution.
- T2*-weighted blood oxygen level-dependent (BOLD) images using a gradient-echo echoplanar imaging sequence were collected with TR=2000ms, TE=27ms, matrix size=64 x 64, voxel size=3.0mm^3^ isotropic, and FA=70 degrees.
- Diffusion weighted data were acquired with a multi-shell diffusion sequence with b-values of 1000 s/mm^2^ with 60 directions each with TR=5520ms, TE=89.5ms, matrix size=210 x 180, voxel size=2mm^3^ isotropic, and FA=90 degrees.
