## supplement_03_dlbs for "CREB: Consistent Reference External Batch Harmonization"

**Dallas Life Brain Study (DLBS) Detailed Description**

Study Design and Participants

The Dallas Life Brain Study (DLBS)^1^ was designed to explore the factors influencing the preservation and decline of cognitive function at various stages with a focus on progression of healthy aging to Alzheimer’s Disease. This was a longitudinal study where participants (21-89 years old) were recruited into three waves 3.5-5 years apart with wave 1 recruiting 464 participants, wave 2 recruiting 338 participants, and wave 3 recruiting 224 participants. All participants were recruited from the Dallas-Fort Worth area.

Inclusion criteria were designed to ensure that participants were relatively cognitively healthy. Participants were excluded if they had suffered from alcoholism or drug addiction in the last 5 years, major heart attack or cancer, psychiatric or neurological disorders, chronic substance abuse, or had been unconscious for more than 10 minutes at some point in their life. Participants were right-handed with a Mini-Mental State Examination ≥26. Participants signed up to three consents: central study inclusive of all cognitive, behavioral, and MRI-based assessments, as well as separate consent forms for the Amyloid and Tau imaging processes. The study was approved by local Institutional Review Boards. All participants gave written informed consent prior to participating.

MRI Acquisition

All scanning was done at University of Texas at Southwestern Medical Center on a 3T Philips Achieva scanner with an 8-channel head coil. Participants underwent MR scanning including T1-weighted MPRAGE, T2-weighted FLAIR, diffusion weighted imaging, arterial spin labeling, and fMRI including Subsequent Memory Task, Semantic Judgement Task, Face/Place Passive Viewing Task, and eyes open resting-state. Participants also underwent Positron Emission Tomography using 18F-AV-45 for amyloid and 18F-AV-1451 for tau, but this data was not used for this analysis.

- T1-weighted magnetization prepared rapid gradient echo (MPRAGE) were collected with repetition time (TR)=8.1ms, echo time (TE)=3.7ms, flip angle (FA)=12deg, field of view (FOV)=204 x 256, and 1mm^3^ isotropic resolution.
- T2-weighted Fluid-Attenuated Inversion Recovery (FLAIR) were collected with TR=1100ms, TE=125ms, FA=not reported, and 0.45x0.45x2.5mm^3^ resolution.
- T2*-weighted blood oxygen level-dependent (BOLD) images using a gradient-echo echoplanar imaging sequence were collected with TR=2000ms, TE=25.0ms, matrix size=64 x 64, voxel size=3.4mm^3^ isotropic, FA=80 degrees, and SENSE acceleration of 2.
- Diffusion weighted data were acquired with a multi-shell diffusion sequence with b-values of 1000 s/mm^2^ with 30 directions with TR=4410ms, TE=51ms, matrix size=224 x 224, voxel size=1.75 x 1.75 x 2mm^3^, and FA=90 degrees.
