## supplement_04_sald for "CREB: Consistent Reference External Batch Harmonization"

**Southwest University adult lifespan dataset (SALD) Detailed Description**

Study Design and Participants

The Southwest University adult lifespan dataset (SALD)^1^ was designed to study the developmental trajectory of the human brain and to understand the functional network changes as a result of aging. This was a large cross-sectional study on 494 participants aged 18 to 96 years old. Young and middle age participants were recruited mostly from students and staff of Southwest University, and older participants were recruited from neighborhood near the University.

The exclusion criteria were MR related exclusion, current psychiatric or neurological disorders, use of psychiatric medications within three months prior to scanning, pregnancy, or history of head trauma. Each participant gave written informed consent prior to participating. The dataset collection was approved by the Research Ethics Committee of the Brain Imaging Center of Southwest University, in accordance with the Declaration of Helsinki.

MRI Acquisition

All scanning was conducted at the Southwest University Center for Brain Imaging using a 3T Siemens Trio – the head coil used was not reported nor was it in the header information. Participants underwent structural T1-weighted scans and 8 minutes of eyes-closed resting state fMRI. Some participants also underwent task based imaging, but this data was not provided publicly.

- T1-weighted magnetization prepared rapid gradient echo (MPRAGE) were collected with repetition time (TR)=1900ms, echo time (TE)=2.52ms, flip angle (FA)=90deg, field of view (FOV)=256 x 256, and 1mm^3^ isotropic resolution.
- T2*-weighted blood oxygen level-dependent (BOLD) images using a gradient-echo echoplanar imaging sequence were collected with TR=2000ms, TE=30ms, matrix size=220 x 220, voxel size=3.4 x 3.4 x 4 mm^3^, and FA=90 degrees.
