## supplement_05_bilingual for "CREB: Consistent Reference External Batch Harmonization"

**Bilingualism and the brain (Bilingual) Detailed Description**

Study Design and Participants

The Bilingualism and the brain (Bilingual)^1^ study designed to examine the neural correlates of bilingualism in adults. They recruited 65 healthy adults between 18 to 52 years old who spoke a variety of first languages, but all spoke English as their second language.

There were minimal exclusion criteria for the study though these are not reported in detail in the original manuscript. All participants were living in the United Kingdom, and all spoke English as a second language and were presumably able to go into the MR scanner. All participants completed an English proficiency test of the Oxford Quick Placement test, and all scored at least high-intermediate. The research procedures in this study were approved by the University of Reading Research Ethics Committee. All participants gave written informed consent and confirmed no contraindication to MRI scanning.

MRI Acquisition

All scanning was performed on a 3T Siemens Magnetom Prisma scanner with a 32-channel head coil. Participants underwent T1-weighted structural scans, a 10-minute resting state fMRI with eyes open, and a diffusion weighted scan.

- T1-weighted magnetization prepared rapid gradient echo (MPRAGE) were collected with repetition time (TR)=2400ms, echo time (TE)=2.41ms, flip angle (FA)=8deg, field of view (FOV)=246 x 256, and 0.7mm^3^ isotropic resolution.
- T2*-weighted blood oxygen level-dependent (BOLD) images using a gradient-echo echoplanar imaging sequence were collected with TR=1500ms, TE=30ms, matrix size=192 x 192, voxel size=2.1 x 2.1 x 2.0mm^3^, and FA=66 degrees.
- Diffusion weighted data were acquired with a multi-shell diffusion sequence with b-values of 1000 s/mm^2^ with 64 directions each with TR=1800ms, TE=70ms, matrix size=128 x 128, voxel size=2mm^3^ isotropic, and FA=90 degrees.
