## supplement_06_ucla_cnp for "CREB: Consistent Reference External Batch Harmonization"

**UCLA Consortium for Neuropsychiatric Phenomics LA5c Study (UCLA-CNP) Detailed Description**

Study Design and Participants

The UCLA Consortium for Neuropsychiatric Phenomics LA5c Study (UCLA-CNP)^1^ was designed to study the dimensional structure of memory and cognitive function in healthy individuals and those with neuropsychiatric disorders. They recruited healthy control participants (n=138) as well as individuals with schizophrenia (n=58), bipolar disorder (n=49), and attention-deficit/hyperactivity disorder (n=45). All participants were aged 21-50 years and recruited through community advertisements from the Los Angeles area. For this analysis, we only used data from healthy control participants.

Inclusion criteria were 21-50 years old and any racial/ethnic category with primary language either English or Spanish; 8 years education, no significant medical illness, visual acuity 20/60 minimum; urinalysis negative for drugs including Cocaine; Methamphetamine; Morphine; THC; and Benzodiazepines. Participants in the healthy group were excluded if they had lifetime diagnoses of Schizophrenia or Other Psychotic Disorder, Bipolar I or II Disorder, or Substance Abuse or Dependence (not counting caffeine or nicotine); or current Major Depressive Disorder; suicidality; Anxiety Disorder (Obsessive Compulsive Disorder, Panic Disorder, Generalized Anxiety Disorder, Post-Traumatic Stress Disorder), and attention deficit hyperactivity disorder (ADHD). ADHD criteria were assessed using Adult ADHD Interview; healthy participants were screened for sub-threshold ADHD, defined as 4 or more ADHD inattentive or hyperactive/impulsive symptoms in either childhood or adulthood; in addition, they could not have had medication treatment for ADHD within the prior 12 months. Each of the patient groups (Schizophrenia, Bipolar Disorder, and ADHD) excluded anyone with one of these other diagnoses; stable medications were permitted for the patients. For MRI studies we excluded participants who were left-handed, who believed they might be pregnant, or had other contraindications to scanning (e.g., claustrophobia, metal in body, body too large to fit in scanner).

MRI Acquisition

Scanning was done on one of two 3T Siemens Trio scanners located at the Ahmanson-Lovelace Brain Mapping Center or the Staglin Center for Cognitive Neuroscience with a Siemens HeadMatrix coil though the number of channels were not reported. Participants underwent T1-weighted structural scans, diffusion weighted scans, 5-minute resting state fMRI, as well as multiple fMRI tasks including Balloon Analog Risk Task, Paired-associate Memory Task, Breath Hold Task, Stop Signal Task, Spatial Capacity working memory task (SCAP), and Task Switching Task.

- T1-weighted magnetization prepared rapid gradient echo (MPRAGE) were collected with repetition time (TR)=1900ms, echo time (TE)=2.26ms, flip angle (FA)=7deg, field of view (FOV)=256 x 256, and 1mm^3^ isotropic resolution.
- T2*-weighted blood oxygen level-dependent (BOLD) images using a gradient-echo echoplanar imaging sequence were collected with TR=2000ms, TE=30ms, matrix size=64 x 64, voxel size=3 x 3 x 4mm^3^, and FA=90 degrees.
- Diffusion weighted data were acquired with a multi-shell diffusion sequence with b-values of 1000 s/mm^2^ with 64 directions each with TR=9000ms, TE=93ms, matrix size=96 x 96, voxel size=2mm^3^ isotropic, and FA=90 degrees.
