## supplement_07_music for "CREB: Consistent Reference External Batch Harmonization"

**Functional connectivity of music induced analgesia in fibromyalgia (Music) Detailed Description**

Study Design and Participants

The Functional connectivity of music induced analgesia in fibromyalgia^1^ was a dataset focused on studying the biological mechanism of music induced analgesia in patient who experience pain from fibromyalgia (FM). This was a cross-sectional study that recruited 40 participants (20 FM and 20 matched healthy control) aged 21–70years old. FM participants were recruited through a fibromyalgia support group and from the General Hospital of the Health Government Department both located in Queretaro, Mexico; Healthy control participants were recruited using flyers placed in the Instituto de Neurobiología. They recruited all women for this study given the difficulties of obtaining male participants with a complete diagnosis of FM.

Inclusion criteria for FM patient were meeting the Fibromyalgia 1990 and 2010 criteria; women with Woman with FM were diagnosed by a trained Rheumatologist; Spontaneous, continuous and intense pain in daily life (visual analog rating scale > 5 average of a month); Right-handed. Inclusion criteria for healthy controls were right-handed healthy adult woman. Exclusion criteria for FM patients and healthy controls were impossibility to move or walk; Uncontrolled endocrine problems; Neurological alterations (i.e. stroke, epilepsy, recent traumatic brain injury); auditory problems; MRI contraindications (i.e., metal prosthetics); Pregnancy and/or breast-feeding. Elimination Criteria for FM patients and healthy control were excessive artifacts in MRI or probable pathological findings in MRI. The study was conducted in accordance with the Declaration of Helsinki and approved by Bioethics Committee of the Instituto de Neurobiología, Universidad Nacional Autónoma de México. All participants gave written informed consent before the study.

MRI Acquisition

All scanning was done at Instituto de Neurobiología of the Universidad Nacional Autónoma de México using a 3T GE Discovery MR750 scanner with commercial 32-channel head coil. FM patient group were instructed to not take painkillers on the day of testing, and HC group were assured that they did not experience any type of pain on the day of testing. Participants were not given any instructions about the music or pink noise. For each participant, 2 five minute runs of rs-FMRI were taken before and after 5 minutes of music and noise session. They were instructed to keep their eyes open in the scanner without thinking anything in particular. Each participant has 3 modalities of images: T1 weighted anatomical imaging, diffusion weighted tensor imaging, and 4 resting state bold images. For this analysis, we used anatomical and resting state bold images.

- T1-weighted anatomical images using the FSPGR BRAVO pulse were collected with repetition time (TR)=7.7ms, echo time (TE)=3.2ms, flip angle (FA)=12deg, field of view (FOV)=256 x 256, and 1.1 x 1 x 1 mm3 resolution.
- T2*-weighted blood oxygen level-dependent (BOLD) images using a gradient-echo echoplanar imaging sequence were collected with TR=3000ms, TE=40ms, matrix size=128 x 128, voxel size=2 x 2 x 3mm^3^ resolution, and FA=90 degrees.
- Diffusion weighted data were acquired with a multi-shell diffusion sequence with b-values of 1000 s/mm^2^ with 60 directions with TR=7000ms, TE=80.8ms, matrix size=256 x 256, voxel size=1 x 1 x 2mm^3^ resolution, and FA=90 degrees.
