## supplement_08_placebo for "CREB: Consistent Reference External Batch Harmonization"

**Brain connectivity predicts placebo response across chronic pain clinical trials (Placebo) Detailed Description**

Study Design and Participants

The Brain connectivity predicts placebo response across chronic pain clinical trials (Placebo) dataset^1^ was focused on identifying whether clinical placebo response can be predicted from patient’s brain connectivity from resting state fMRI. The hypothesis of the study was that functional connectivity could be used as functional markers for analgesia associated with placebo treatment. This was a cross-sectional study that recruited 76 participants (56 Chronic knee osteoarthritis (OA) pain patients and 20 healthy controls) aged 44-78 years through public advertisements and Northwestern University-affiliated clinics.

Inclusion criteria for healthy controls were age 45-80 years old and ability to give informed consent. Additional inclusion criteria for OA patients were VAS pain score >5/10 within 48 hours of the phone screen and visit 1 (Screening); knee OA for a minimum of 12 months; need for daily pain medication to manage symptoms of OA; satisfies American College of Rheumatology (ACR) criteria for OA including Kellgren-Lawrence radiographic OA grades II-IV. Exclusion criteria were currently taking MAO inhibitors or any centrally acting medication for analgesia or depression; narrow angle glaucoma; uncontrolled hypertension; co-existing inflammatory arthritis, fibromyalgia or other chronic pain; pregnant or trying to become pregnant or lactating; depression; substantial alcohol use or history of liver disease; use of triptans, serotonin precursors (tryptophan), CYP1A2 inhibitors, thioridazine, and antidepressants; diabetes; conditions that investigators felt may interfere with following study instructions or put participant at risk; and MR safety exclusions. The study was approved by Northwestern University Institutional Review Board. All participants gave written consent prior to participating.

MRI Acquisition

All scanning was done at Northwestern University using a 3T Siemens Trio whole body scanner with echo planar imaging (EPI) capability with a head-coil, but the number of channels were not reported. T1-weighted structural images and resting state fMRI were acquired for each participant on the same day. In our analysis, we used both the structural and resting state fMRI.

- T1 weighted magnetization prepared rapid gradient echo (MPRAGE) were collected with repetition time (TR) = 2.5ms, echo time (TE)=3.36ms, flip angle (FA) = 9 degrees, field of view (FOV) = 256 × 256, and 1mm^3^ isotropic resolution.
- T2*-weighted blood oxygen level-dependent (BOLD) images using a gradient-echo echoplanar imaging sequence were collected with TR=2500ms, TE=30ms, matrix size=64×64, voxel size= 3.4 × 3.4 × 3 mm, FA= 90 degrees.
