## supplement_09_depression for "CREB: Consistent Reference External Batch Harmonization"

**Resting State with closed eyes for patients with depression and healthy participants (Depression)** **Detailed Description**

Study Design and Participants

The Resting State with closed eyes for patients with depression and healthy participants (Depression) dataset^1,2^ was designed to characterize within network and between network functional connectivity differences in patients with mild to moderate depression. The dataset was designed to monitor changes in functional connectivity of depressed participants as they received nonpharmacological treatments to see if baseline connectivity could predict treatment response, and if connectivity changes correlates with clinical improvements. This was a cross-sectional study that recruited 72 participants (21 healthy, 51 depressed) aged 19-52 years.

Inclusion criteria were healthy or depressed participants able to give informed consent. Exclusion criteria were neurological or psychotic level mental disorders; psychotropic medication or drugs severely influencing blood flow; contraindications to MRI; depression condition that is bipolar, seasonal, or secondary to other disease; IQ $\leq$70 with Raven Progressive Matrices test. This study’s protocol was in accordance with Helsinki Declaration and was approved by local ethic board of Institute of Molecular Biology and Biophysics in Novosibirsk, Russia. All participants signed informed consent prior to inclusion in study.

MRI Acquisition

All scanning was done at International Tomography Center, Novosibirsk, using a 3T Philips Ingenia scanner. For each participant, T1-weighted structural and a 6-minute resting state fMRI were collected with eyes closed. Participants were instructed to lie still with eyes closed for 6 minutes for resting state fMRI. In analysis, we used both the structural and resting state bold images from all healthy control participants.

- T1-weighted 3D turbo field echo (3D T1 TFE) was collected with repetition time (TR)=not reported, echo time (TE) = not reported, flip angle (FA) =not reported, field of view (FOV)=228×228, and voxel resolution = 0.87 × 0.87 × 1.0 mm.
- T2*-weighted blood oxygen level-dependent (BOLD) images using a gradient-echo echoplanar imaging sequence were acquired with TR=2500ms, TE=35ms, matrix size=112×112, voxel size=2 × 2 × 5 mm, FA=90 degrees.
