## supplement_10_camcan for "CREB: Consistent Reference External Batch Harmonization"

**The Cambridge Centre for Ageing and Neuroscience (Cam-CAN) Detailed Description**

Study Design and Participants

The Cambridge Centre for Ageing and Neuroscience (Cam-CAN)^1^ study was designed to characterize age-related changes in cognition and brain structure and function, and to uncover the neurocognitive mechanisms that support healthy cognitive aging. Cam-CAN was a large cross-sectional study of 700 participants aged 18-87 years old who were drawn from a larger population-based sample (n=3000). Participants were recruited from the Cambridge City area.

Participants had to have a mini mental state examination above 24, no MRI or MEG contraindications, not have other conditions including serious head injury, current drug abuse, or current serious psychiatric condition, not have poor hearing ability, and must have adequate English. This study was conducted in compliance with the Helsinki Declaration, and was approved by the local ethics committee, Cambridgeshire 2 Research Ethics Committee. All participants gave written informed consent prior to participating.

MRI Acquisition

All MR scan was conducted at a single site (MRC-CBSU) with T Siemens TIM Trio scanner with a 32-channel head coil. MR scans included a T1-weighted and a T2-weighted structural, diffusion weighted imaging, magnetization transfer imaging, and functional imaging including an ~8.5-minute eyes closed resting state fMRI and a movie watching task and sensorimotor task.

- T1-weighted magnetization prepared rapid gradient echo (MPRAGE) were collected with repetition time (TR)=2250ms, echo time (TE)=2.99ms, flip angle (FA)=9deg, field of view (FOV)=256 x 240, and 1mm^3^ isotropic resolution.
- T2-weighted Sampling Perfection with Application optimized Contrasts using different flip angle Evolution (SPACE) were collected with TR=2800ms, TE=408ms, FA=92 degrees, FOV=256 x 256, and 1mm^3^ isotropic resolution.
- T2*-weighted blood oxygen level-dependent (BOLD) images using a gradient-echo echoplanar imaging sequence were collected with TR=1970ms, TE=30ms, matrix size=192 x 192, voxel size=3 x 3 x 4.4mm^3^, and FA=78 degrees.
- T2*-weighted blood oxygen level-dependent (BOLD) images for the movie watching task using a gradient-echo echoplanar imaging sequence were collected with TR=2470ms, multi-echo TE=9.4, 21.2, 33, 45, and 57ms, matrix size=192 x 192, voxel size=3 x 3 x 4.4mm^3^, and FA=78 degrees.
- Diffusion weighted data were acquired with a multi-shell diffusion sequence with b-values of 1000 and 2000 s/mm^2^ with 30 directions each with TR=9100ms, TE=104ms, matrix size=192 x 192, voxel size=2mm^3^ isotropic, and FA=not reported.
