## supplement_11_aging for "CREB: Consistent Reference External Batch Harmonization"

**Neurocognitive aging data release with behavioral, structural, and multi-echo functional MRI measures (Aging) Detailed Description**

Study Design and Participants

The Neurocognitive aging data release with behavioral, structural, and multi-echo functional MRI measures (Aging) dataset^1^ was designed to investigate differences across age cohorts in cognition and brain health, and how they intersect to shape late-life development. It was used to study the pattern of functional network dedifferentiation and other age-related functional changes with age, and how such dedifferentiation and re-organizing of network connectivity could impact cognitive functions. This was a cross-sectional and multi-site study that recruited 301 healthy adult participants aged 18-89+ years from Ithaca, New York and Toronto, Canada. Two age groups, young (age 18-34 years) and old (age 60-89 years) were recruited, the younger group served as reference to measure the dedifferentiation and integration of functional networks in the older group^2^.

Inclusion criteria were right-handed healthy adult participants with normal or corrected-to-normal vision who were able to give inform consent. Exclusion criteria were individuals with a history of neurological or other medical illness known to impact cognition; acute or chronic psychiatric illness; those undergoing current or recent treatment with psychotropic medication; and those having recently experienced significant changes to health status at the time of the eligibility interview; scored below 27/30 on MMSE coupled with scored in the bottom 25th percentile of age-adjusted scores for fluid cognition. Young adults were screen with Beck Depression Inventory^3^ and older adult were screened with Geriatric Depression Scale^4^. The study was in-compliance with IRB at Cornell University and York University, and each participant gave written informed consent prior to participating in the study.

MRI Acquisition

Scanning was conducted from 2 sites with a 3T GE Discovery MR750 and 32-channel head coil at the Cornell Magnetic Resonance Imaging Facility (237 participants scanned here) or on a 3T Siemens TimTrio MRI scanner with a 32-channel head coil at the York University Neuroimaging Center in Toronto (63 participants scanned here). At both sites, participants were asked to stay awake and lie still with their eyes open, breathing, and blinking normally in the darkened scanner bay. Each participant underwent T1-weighted structural scans, and two 10-minute resting state multi-echo fMRI, and a subset of 110 older adults and 148 younger adults have T2-weighted FLAIR scans. All data were collected on the same day scanning day for each participant. For analysis, we used both the T1 and T2-FLAIR (if available) images and the two resting state fMRI.

- T1-weighted magnetization prepared rapid gradient echo (MPRAGE) were collected with repetition time (TR) = 2530ms, echo time (TE) = 3.4ms, field of view (FOV) = 256×256, Flip angle (FA) = 7 degrees, voxel size=1mm^3^ isotropic, at Cornell Magnetic Resonance Imaging Facility. T1-weighted generalized auto calibrating partially parallel acquisition (GRAPPA) were collected with TR=1900ms, TE=2.52ms, FOV = 192×256, FA = 9 degrees, voxel size = 1mm isotropic, in York University Neuroimaging Center in Toronto.
- T2-weighted FLAIR sequences were acquired with TR = 12000ms, TE = 95ms, matrix size=256×256, FA = 160 degrees, voxel size = 1 × 1 × 3mm^3^, at Cornell Magnetic Resonance Imaging Facility. T2-weighted FLAIR sequences were acquired with TR = 12000ms, TE = 95ms, matrix size=384×512, FA = 160 deg, voxel size =0.8 × 0.8 × 3mm^3^, in York University Neuroimaging Center in Toronto. (12 participants have 46 slices due to technician error^5^)
- T2*-weighted blood oxygen level-dependent (BOLD) images were collected with multi-echo (ME) EPI sequence with TR = 3000ms TE_1_ = 13.7ms, TE_2_=30ms, TE_3_=47ms, FA= 83 deg, matrix size = 72 × 72, field of view (FOV)= 210mm, voxel size = 3mm^3^ isotropic at Cornell Magnetic Resonance Imaging Facility; and with TR = 3000ms, TE_1_ = 14 ms, TE_2_=29.96 ms, TE_3_=45.92ms, FA = 83 degrees, matrix size = 64×64, voxel size = 3.4× 3.4×3mm^3^ at York University Neuroimaging Center in Toronto. (sub-149 has 206 total volumes collected instead of 204^5^)
