## supplement_12_glia for "CREB: Consistent Reference External Batch Harmonization"

**A large-scale dataset of pre- and post-surgical MRI data in patients with chronic trigeminal neuralgia (Glia) Detailed Description**

Study Design and Participants

The pre- and post-surgical MRI data in patients with chronic trigeminal neuralgia (Glia) dataset^1^ focused on characterizing brain connectivity changes after surgery of trigeminal neuralgia. This was a longitudinal study that recruited 160 participants aged 31-84 years (112 Pre-surgical Trigeminal Neuralgia (PTN) participants and 48 healthy controls). 58 people of the PTN patient group have 6 months follow up scans additional to their baseline scans. We only used data from healthy controls.

The detailed inclusion and exclusion criteria for this dataset was not reported on OpenNeuro. PTN inclusion criteria specify participants need to be adults with diagnosed primary trigeminal neuralgia (according to The International Classification of Headache Disorders criteria). Ethics approval for the research protocol was given by local ethic committee of Federal Center of Neurosurgery. All participants provided written informed consent prior to participating the study in accordance with the Declaration of Helsinki.

MRI Acquisition

All scanning was done with 3T Philips Ingenia scanner in Federal Neurosurgical Center, Novosibirsk, Russia. For each participant, they collected data on T1-weighted and T2-weighted structural scans, diffusion weighted scans, and resting state fMRI scans. They did not report whether participants had eyes closed or open. For our analysis, we only used the structural and resting state fMRI from control participants.

- T1-weighted magnetization prepared rapid gradient echo (MPRAGE) were collected with repetition time (TR) = 6.6ms, echo time (TE)=2.96ms, flip angle (FA)=8 deg, field of view(FOV)=340×512, 0.5mm^3^ isotropic resolution.
- T2-weighted 3D turbo spin-echo (TSE) were collected with TR= 3500ms, TE= 300ms, FA= 90 deg, FOV= 360 × 512, and voxel size = 0.5mm^3^ isotropic resolution.
- T2*-weighted blood oxygen level-dependent (BOLD) images using a gradient-echo echoplanar imaging sequence were collected with TR=3000ms, TE=30ms, matrix size = 80×80, voxel size = 3 mm^3^ isotropic, FA=90 degrees. The slice order is **standard Philips interleaved slice acquisition order**, in which slices were acquired in the following sequence ascending starting with first slice every three slices then repeating on slices 2 and 3 once completed.
- Diffusion weighted data were acquired using single-shot spin-echo echo-planar imaging (EPI) sequence with b-value and 1500 s/mm^2^ with 64 directions each with TR = 10490 ms, TE = 111.9 ms, matrix size = 128 × 128, voxel size = 2 mm³ isotropic, and FA = 90 degrees,
