## supplement_13_fmriprep for "CREB: Consistent Reference External Batch Harmonization"

**Detailed Description fMRIprep Processing**

To get grey matter volume measures, we ran fMRIPrep with the flag for performing anatomical workflows only. For datasets with multiple longitudinal sessions, we use a BIDS filter to make sure only anatomical images of the first scan were processed since by default, anatomical images from different sessions are combined together. For datasets that already completed skull stripping, we skipped skull stripping.

**Anatomical workflow**

We ran the anatomical only workflow using fMRIPrep 23.2.3^1,2^ , which is based on Nipype 1.8.6^3^. T1-weighted (T1w) images for each dataset were organized into BIDS format. T1w images were corrected for intensity non-uniformity (INU) with N4BiasFieldCorrection ^4^, distributed with ANTs 2.5.0^5^, and used as T1w-reference throughout the workflow. The T1w-reference was then skull-stripped with a Nipype implementation of the antsBrainExtraction.sh workflow (from ANTs), using OASIS30ANTs as target template. Brain tissue segmentation of cerebrospinal fluid (CSF), white-matter (WM) and gray-matter (GM) was performed on the brain-extracted T1w using FSL Fast^6^. Volume-based spatial normalization to one standard space (MNI152NLin2009cAsym) was performed through nonlinear registration with antsRegistration (ANTs 2.5.0), using brain-extracted versions of both T1w reference and the T1w template. The following templates were selected for spatial normalization and accessed with TemplateFlow 23.1.0^7^: ICBM 152 Nonlinear Asymmetrical template version 2009c ^8^ and TemplateFlow ID of MNI152NLin2009cAsym.

**Functional Workflow**

For each of the BOLD runs found per participant across all tasks and sessions, the following preprocessing was performed. First, a reference volume was generated, using a custom methodology of fMRIPrep, for use in head motion correction. Head-motion parameters with respect to the BOLD reference (transformation matrices, and six corresponding rotation and translation parameters) were estimated before any spatiotemporal filtering using mcflirt^9^. The BOLD reference was then co-registered to the T1w reference using mri_coreg (FreeSurfer) followed by flirt^10^ with the boundary-based registration ^11^ cost-function. Co-registration was configured with six degrees of freedom. Several confounding time-series were calculated based on the preprocessed BOLD: framewise displacement (FD), DVARS and three region-wise global signals. FD was computed using two formulations following Power et al^12^ absolute sum of relative motions and Jenkinson’s relative root mean square displacement between affines^9^. FD and DVARS were calculated for each functional run, both using their implementations in Nipype (following the definitions by Power et al. 2014^12^). The three global signals were extracted within the CSF, the WM, and the whole-brain masks. Additionally, a set of physiological regressors were extracted to allow for component-based noise correction (CompCor from Behzadi et al^13^). Principal components were estimated after high-pass filtering the preprocessed BOLD time-series (using a discrete cosine filter with 128s cut-off) for the two CompCor variants: temporal (tCompCor) and anatomical (aCompCor). tCompCor components were then calculated from the top 2% variable voxels within the brain mask. For aCompCor, three probabilistic masks (CSF, WM and combined CSF+WM) were generated in anatomical space. The implementation differs from that of Behzadi et al. instead of eroding the masks by 2 pixels on BOLD space, a mask of pixels that likely contain a volume fraction of GM is subtracted from the aCompCor masks. This mask is obtained by thresholding the corresponding partial volume map at 0.05, and it ensures components were not extracted from voxels containing a minimal fraction of GM. Finally, these masks were resampled into BOLD space and binarized by thresholding at 0.99 (as in the original implementation). Components were also calculated separately within the WM and CSF masks. For each CompCor decomposition, the k components with the largest singular values were retained, such that the retained components’ time series were sufficient to explain 50 percent of variance across the nuisance mask (CSF, WM, combined, or temporal). The remaining components were dropped from consideration. The head-motion estimates calculated in the correction step were also placed within the corresponding confounds file. The confound time series derived from head motion estimates and global signals were expanded with the inclusion of temporal derivatives and quadratic terms for each^14^. Frames that exceeded a threshold of 0.5 mm FD or 1.5 standardized DVARS were annotated as motion outliers. Additional nuisance timeseries were calculated by means of principal components analysis of the signal found within a thin band (crown) of voxels around the edge of the brain, as proposed by^15^. All resamplings can be performed with a single interpolation step by composing all the pertinent transformations (i.e. head-motion transform matrices, susceptibility distortion correction when available, and co-registrations to anatomical and output spaces). Gridded (volumetric) resamplings were performed using nitransforms, configured with cubic B-spline interpolation.
