## supplement_14_xcpd for "CREB: Consistent Reference External Batch Harmonization"

**Detailed Description XCPD Processing**

We use eXtensible Connectivity Pipeline-DCAN (XCP-D) to post-process the outputs of fMRIPrep version 23.2.3^1–3^. The following atlases were used in the workflow: the Schaefer Supplemented with Subcortical Structures (4S) atlas ^4–8^. The atlas was warped to the same space and resolution as the BOLD file. For anatomical data, native-space T1w images were transformed to MNI152NLin2009cAsym space at 1 mm^3^ resolution. For functional data, we use each run found per participants across all tasks and sessions and with following post-processing procedure described below.

Non-steady-state volumes were extracted from the preprocessed confounds and were discarded from both the BOLD data and nuisance regressors. Framewise displacement was calculated from the motion parameters using the formula from Power et al.^9^ with a head radius of 50 mm. Volumes with framewise displacement greater than 0.5 mm were flagged as high-motion outliers for the sake of later censoring ^9^. Nuisance regressors were selected according to the “acompcor” strategy. The top 5 aCompCor principal components from the white matter and cerebrospinal fluid compartments were selected as nuisance regressors ^10^ along with the six motion parameters and their temporal derivatives ^11,12^. As the aCompCor regressors were generated on high-pass filtered data, the associated cosine basis regressors were included. This has the effect of high-pass filtering the data as well. Nuisance regressors were regressed from the BOLD data using a denoising method based on Nilearn*’s* approach. Any volumes censored earlier in the workflow were first cubic spline interpolated in the BOLD data. Outlier volumes at the beginning or end of the time series were replaced with the closest low-motion volume’s values, as cubic spline interpolation can produce extreme extrapolations. The timeseries were low-pass filtered using a second-order Butterworth filter, in order to retain signals below 0.15 Hz. The same filter was applied to the confounds. The resulting time series were then denoised via linear regression, in which the low-motion volumes from the BOLD time series and confounds were used to calculate parameter estimates, and then the interpolated time series were denoised using the low-motion parameter estimates. The interpolated time series were then censored using the temporal mask. The denoised BOLD was smoothed using Nilearn with a Gaussian kernel (FWHM=8.0 mm). Processed functional timeseries were extracted from the residual BOLD signal with Nilearn’s NiftiLabelsMasker for the atlases. Corresponding pair-wise functional connectivity between all regions was computed for each atlas, which was operationalized as the Pearson’s correlation of each parcel’s unsmoothed timeseries. In cases of partial coverage, uncovered voxels (values of all zeros or NaNs) were either ignored (when the parcel had >0.0% coverage) or were set to zero (when the parcel had <0.0% coverage).
