## supplement_15_creb_augmentation for "CREB: Consistent Reference External Batch Harmonization"

**fMRI Data Augmentation Detailed Description**

Data augmentation increases the number of connectivity matrices estimated from each run of each participant. We do this by calculating the connectivity on a subset of the timeseries. Augmentation was done 4 times for each run by selecting a random starting point and random duration that has minimum of 3 min and maximum base on the total length of the run. We calculated the length of the subarray that we should select using the duration in minutes of each run. The resulting augmented correlation matrices will have suffix with starting point volume index, duration in number of volumes, and duration in seconds. The table below of the TR for each task from each study and the duration min, mean, and max we used for augmentation.

| **Dataset Name** | **Task/Type** | **TR (s)** | **Min Duration (min)** | **Mean Duration (min)** | **Max Duration (min)** |
| --- | --- | --- | --- | --- | --- |
| **HCP-YA** | **rest** | 0.72 | 3 | 10 | 14 |
| **DLBS** | **rest** | 2 | 3 | 4 | 5.5 |
|  | **Scenes** | 2 | 3 | 4 | 5.5 |
|  | **VentralVisual** | 2 | 3 | 5 | 6.5 |
|  | **Words** | 2 | 3 | 5.5 | 7.5 |
|  | **Hypercapnia** | 2 | 3 | 5.5 | 7 |
| **BVDCC** | **rest** | 2 | 3 | 7 | 9 |
|  | **nbk** | 2 | 3 | 7 | 8.5 |
|  | **tsw** | 2 | 3 | 5.5 | 7 |
|  | **gng** | 2 | 3 | 5.5 | 7 |
| **Depression** | **rest** | 2.5 | 3 | 3.5 | 4 |
| **Music** | **rest** | 3 | 3 | 4 | 5 |
| **UCLA-CNP** | **rest** | 2 | 3 | 4 | 5 |
|  | **bart** | 2 | 3 | 7 | 8.5 |
|  | **scap** | 2 | 3 | 7 | 9.5 |
|  | **stopsignal** | 2 | 3 | 5 | 6 |
|  | **taskswitch** | 2 | 3 | 5 | 6.5 |
|  | **pamenc** | 2 | 3 | 6 | 8 |
|  | **pamret** | 2 | 3 | 7 | 8.5 |
| **Placebo** | **rest** | 2.5 | 3 | 9 | 12 |
| **SALD** | **rest** | 2 | 3 | 6 | 8 |
| **Bilingual** | **rest** | 1.5 | 3 | 5 | 7 |
| **Aging** | **rest** | 3 | 3 | 7 | 10 |
| **CamCAN** | **rest** | 1.97 | 3 | 5 | 8 |
|  | **movie** | 2.47 | 3 | 5 | 8 |
| **Glia** | **rest** | 3 | 3 | 7 | 9 |
