## supplement_16_creb_imputation for "CREB: Consistent Reference External Batch Harmonization"

**fMRI Connectivity Data Imputation Detailed Description**

There were missing values for various regions in various studies either due to coverage or low signal-to-noise ratio or some other reason. In addition, our choice of the XCP-D’s built in 4S456 atlas was relatively more refined compared to other commonly used atlases like AAL atlas or Power atlas, some parcels had missing values. Common regions that have missing values were orbital frontal cortex (OFC) due to magnetic susceptibility artifact. For each study, we report the total runs, number of runs with at least one missing value, average number of regions of interest (ROIs) missing, average percent of ROIs missing for each study, and range of missing regions.

We conduct a form of imputation where we estimate the mean and variance of the connectivity matrix ignoring missing values and then use that distribution to estimate 3000 random connectivity matrices. We sample from those 3000 matrices when we needed to impute a value. We first assigned correlation matrices from all datasets into train, validation, and test set. For the 8700 correlation matrices (after augmentation) in the training rest folder, we calculate the mean and variance for each edge. We then generated 3000 synesthetic matrices using the mean and variance matrix calculated from the train. Critically, these are only ‘sampled’ from the training set to further prevent data leakage. We then iterated through all matrices in train, validation, test sets, and in external test sets, and imputed missing edges from randomly selected synthetic matrices.

| Dataset | Total Files (N) | Files with Missing ROIs (N (%)) | Average Missing ROIs (mean±std) | Average % Missing (per file) | Range (Missing ROIs) |
| --- | --- | --- | --- | --- | --- |
| Bilingual | 176 | 12 (6.8%) | 0.07±0.25 | 0.01% | [0, 1] |
| BVDCC | 421 | 156 (37.1%) | 0.68±1.16 | 0.15% | [0, 7] |
| Depression | 56 | 52 (92.9%) | 5.79±3.67 | 1.27% | [0, 15] |
| DLBS | 3687 | 2227 (60.4%) | 1.24±3.35 | 0.27% | [0, 67] |
| HCP-YA | 2818 | 26 (0.9%) | 0.56±10.04 | 0.12% | [0, 278] |
| Music | 203 | 134 (66.0%) | 2.06±2.17 | 0.45% | [0, 7] |
| Placebo | 48 | 25 (52.1%) | 0.90±1.00 | 0.20% | [0, 3] |
| SALD | 1009 | 675 (66.9%) | 1.61±2.12 | 0.35% | [0, 21] |
| UCLA-CNP | 282 | 249 (88.3%) | 3.24±2.46 | 0.71% | [0, 10] |
